# Capicua regulates neural stem cell proliferation and lineage specification through control of Ets factors

**DOI:** 10.1101/335984

**Authors:** Shiekh Tanveer Ahmad, Alexandra D. Rogers, Myra J. Chen, Rajiv Dixit, Lata Adnani, Luke S. Frankiw, Samuel O. Lawn, Michael D. Blough, Mana Alshehri, Wei Wu, Stephen M. Robbins, J. Gregory Cairncross, Carol Schuurmans, Jennifer A. Chan

## Abstract

Capicua (Cic) is a transcriptional repressor mutated in the brain cancer oligodendroglioma. Despite its cancer link, little is known of Cic’s function in the brain. Here, we investigated the relationship between Cic expression and cell type specification in the brain. Cic is strongly expressed in astrocytic and neuronal lineage cells but is more weakly expressed in stem cells and oligodendroglial lineage cells. Using a new conditional *Cic* knockout mouse, we show that forebrain-specific *Cic* deletion increases proliferation and self-renewal of neural stem cells. Furthermore, *Cic* loss biases neural stem cells toward glial lineage selection, expanding the pool of oligodendrocyte precursor cells (OPCs). These proliferation and lineage selection effects in the developing brain are dependent on de-repression of Ets transcription factors. In patient-derived oligodendroglioma cells, CIC re-expression or ETV5 blockade decreases lineage bias, proliferation, self-renewal and tumorigenicity. Our results identify Cic is an important regulator of cell fate in neurodevelopment and oligodendroglioma, and suggest that its loss contributes to oligodendroglioma by promoting proliferation and an OPC-like identity via Ets overactivity.

## INTRODUCTION

The identification of genes recurrently mutated in cancer often presents opportunities to uncover previously unappreciated mechanisms regulating normal development, and vice versa. The transcriptional repressor Capicua (*CIC*) has been identified as a likely tumour suppressor gene, as recurrent mutations in *CIC* and/or reduced expression of CIC are found in several cancer types. In the brain, *CIC* mutations are nearly exclusively found in oligodendrogliomas (ODGs) – glial tumors that are composed of cells resembling oligodendrocyte precursor cells (1, 2). Indeed, concurrent *IDH1/2* mutation, single-copy whole-arm losses of 1p and 19q, and mutation of the remaining copy of *CIC* on chr 19q13 are highly characteristic of ODG and are not found in other cancer types (3–5). These associations suggest a unique relationship between CIC and glial biology.

Prior work in Drosophila and in mammalian cultured cells has shown that Cic is a transcriptional repressor downstream of receptor tyrosine kinase (RTK) signalling (6). Binding of Cic to the sequence T(G/C)AATG(G/A)A in enhancers and promoters leads to transcriptional repression of its target genes (7, 8). This default repression is relieved upon RTK signalling (6, 9–11), permitting transcription of targets – among which are *PEA3/ETS* transcription factors *ETV1/4/5* (12). To date, there is limited knowledge of Cic’s function in mammalian development or in the brain. Recently, Yang et al. using *Cic* conditional knockout mice, reported that Cic loss increases a population of proliferating Olig2+ cells in the brain, and that) loss of Cic potentiates glial tumorigenesis in a glioma model driven by *PDGFR* (13). The mechanisms by which Cic loss caused those findings, however, remained undefined. Meanwhile, in the non-neoplastic context, Lu et al. recently showed that impairing the interaction of the Cic with the protein Ataxin 1 results in spectrum of neurobehavioral and neurocognitive phenotypes, as well as abnormal maturation and maintenance of populations of cortical neurons (14) – indicating a role for Cic in neuronal biology as well. Deciphering Cic’s mechanisms of action as a neurodevelopmental regulator may shed light on these varied phenotypes resulting from Cic loss or abnormal function.

Here, we examine the temporal and spatial pattern of Cic expression in the mammalian cerebral cortex and white matter, and take a loss-of-function approach to determine its role in neuronal-glial identity determination. Our results reveal an important role for Cic in regulating the proliferation and lineage specification of neural stem cells during development – with loss favoring NSC proliferation and glial production at the expense of neuronal production. Furthermore, we show that these effects are mediated largely through Cic’s regulation of Ets factors. The proliferative dysregulation is recapitulated in oligodendroglioma cells, where CIC re-expression or blockade of Ets activity reduces tumorigenicity. Our findings reveal an important role for CIC in development and oligodendroglioma, and suggest that Etv5 may be a potential therapeutic target.

## RESULTS

### Nuclear Cic levels are cell type- and stage-specific

As an initial step to investigating Cic’s potential function in forebrain development and oligodendroglioma, we examined its expression in several regions and cell types in the mouse brain. Immunofluorescence staining for CIC and cell-type specific markers revealed that all cell types examined have some detectable level of Cic protein, whether cytoplasmic or nuclear. Because of Cic’s previously identified role as a transcriptional repressor, however, we were particularly interested in the nuclear levels and quantitated Cic nuclear staining intensity across cell types and stages.

Focusing first on the stem cell compartment, we assessed tissue from the dorsal telencephalon (the anlage of the neocortex) at embryonic day (E) E12. At this early neurogenic stage, non-lineage restricted neural stem cells (NSCs) are found in the ventricular zone (VZ) where can be identified by their spatial location and their expression of the transcription factor Sox2 (15). In embryonic Sox2+ VZ cells, Cic was localized predominantly in the cytoplasm, with relatively weak nuclear expression (Figure 1A). This distribution is consistent with the previous demonstration that growth factor/RTK signaling is locally elevated in the embryonic VZ (16, 17) and that Cic nuclear export is regulated by ERK-mediated Cic phosphorylation (6, 9–11). Pools of stem/progenitor cells are not limited to development, but also persist postnatally in the subventricular zone (SVZ) of the lateral ventricle and in the subgranular layer (SGL) of the hippocampal dentate gyrus (18, 19). In the SVZ and SGL of postnatal day (P) P21 and P56 mice, Cic was also weakly expressed within the nucleus in the Sox2+ compartment – in contrast to stronger nuclear expression in adjacent differentiated cells (Figure 1B,C). Thus, in both the embryonic and postnatal brain, neural stem cells are characterized by low levels of nuclear Cic.

**FIGURE 1:**
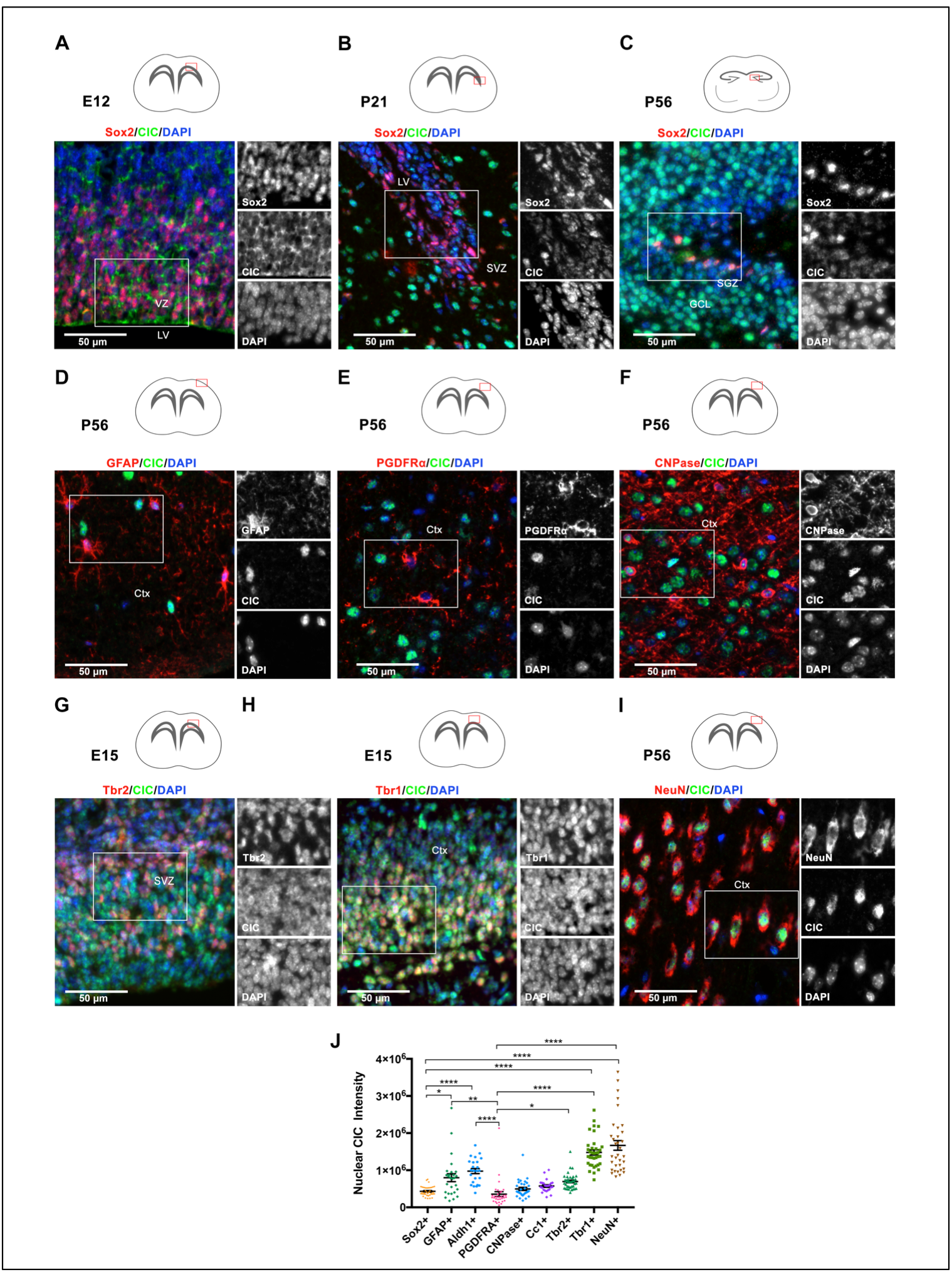
Differential Cic expression among cell types in the developing and mature brain. Representative images of immunofluorescence staining for Cic expression in Sox2+ stem cell populations in (A) E12 subventricular zone, (B) P56 hippocampal dentate gyrus subgranular zone, and (C) P56 subventricular zone. Cic expression in glial cells in the adult brain showing localization in (D) Gfap+ cortical astrocytes, (E) Pdgfra+ white matter OPCs, and (F) CNPase+ mature oligodendrocytes. Cic expression in cortical neuronal populations (G) E15 Tbr2+ early-born neurons in the SVZ and IZ, (H) E15 Tbr1+ late-born neurons in the cortical plate, and (I) adult post-mitotic NeuN+ cortical neurons. (J) Quantitation of Cic nuclear staining intensities, plotted for different brain cell types. Each point represents one cell quantitated. Results show quantitation from a minimum of 24 cells per marker from the boxed regions used for image analysis. All quantitation performed on P56 cortex except Tbr2 and Tbr1 which were analyzed at E15. Pairwise comparisons between cell types performed by ANOVA with Tukey’s posthoc test. Data shown as mean ± SD. *p<0.05, **p<0.01, ****p<0.0001. LV–lateral ventricle, GCL–granule cell layer, SGZ–subgranular zone, SVZ–subventricular zone, Ctx–cortex.

As cells differentiated, Cic was increasingly localized to the nucleus, but with notable differences between the levels detected in neuronal, astrocytic, and oligodendrocytic cell lineages. In the adult (P56) cortex, NeuN+ neurons displayed the strongest nuclear Cic (Figure 1I). The increase in Cic within the neuronal lineage was detectable during the process of embryonic neurogenesis, with a modest increase detected as cells transitioned from Sox2+ stem cells to Tbr2+ neuronal intermediate progenitors, followed by a more marked elevation in Tbr1+ post-mitotic neurons in the intermediate zone and cortical plate – where nuclear Cic levels approached those of mature neurons in the adult cortex (Figure 1G,H,I, J). In GFAP+ or Aldh1+ astrocytes in the cortex and underlying white matter, nuclear Cic levels also increased relative to Sox2+ cells (Figure 1D,J; Supplemental Figure S1A). Among the three major lineages, the lowest levels of nuclear Cic were found in the oligodendroglial lineage. Olig2 is a bHLH transcription factor that is expressed in cells at or before oligodendrocyte specification and continues to be present throughout the stages of oligodendroglial differentiation (20, 21). Within the Olig2+ population, the subset of Pdgfra+ cells indicative of oligodendrocyte precursor cells (OPCs) had the lowest levels of all, with a modest increase in CNPase+ immature oligodendrocytes, and a further increase in mature CC1+ oligodendrocytes (Figure 1E, F, J; and Suppl Fig S1). Overall, the mean integrated nuclear signal intensity for Cic was significantly lower in Sox2+ and Pdgfra+ cells than in NeuN+ cells (p < 0.0001), Tbr1+ cells (p < 0.0001), or Gfap+ cells (p < 0.01) (Figure 1J and Suppl S1).

As both NSCs and OPCs are cell types that are proliferation-competent, this pattern of lower nuclear Cic in NSCs and OPCs, and higher nuclear Cic expression in neurons and astrocytes, raised the possibility that nuclear CIC bound to its target genes may repress proliferation-related genes and/or early oligodendroglial-promoting programs. We tested these hypotheses with a series of loss-of-function studies.

### Cic-deficiency results in increased glial populations and decreased neuronal populations during forebrain development

Cic contains several critical domains including an HMG-box and a C-terminal C1 domain that together bind to DNA, and a C-terminal Gro-L domain that mediates protein-protein interactions (10, 22–25). We generated *Cic* conditional knockout mice in which exons 2-11 of *Cic* gene were flanked by loxP sites, with the floxed region containing all exons encoding the HMG box. Upon introduction of Cre, exons 2-11 are excised and the remaining exons 12-20 are frameshifted, thus ablating the HMG box, C1 domain, and Gro-L domain (Figure 2A). We used these animals for in vivo studies as well as to generate cell lines with which we could further study proliferation and lineage selection in a cell-autonomous manner.

**FIGURE 2:**
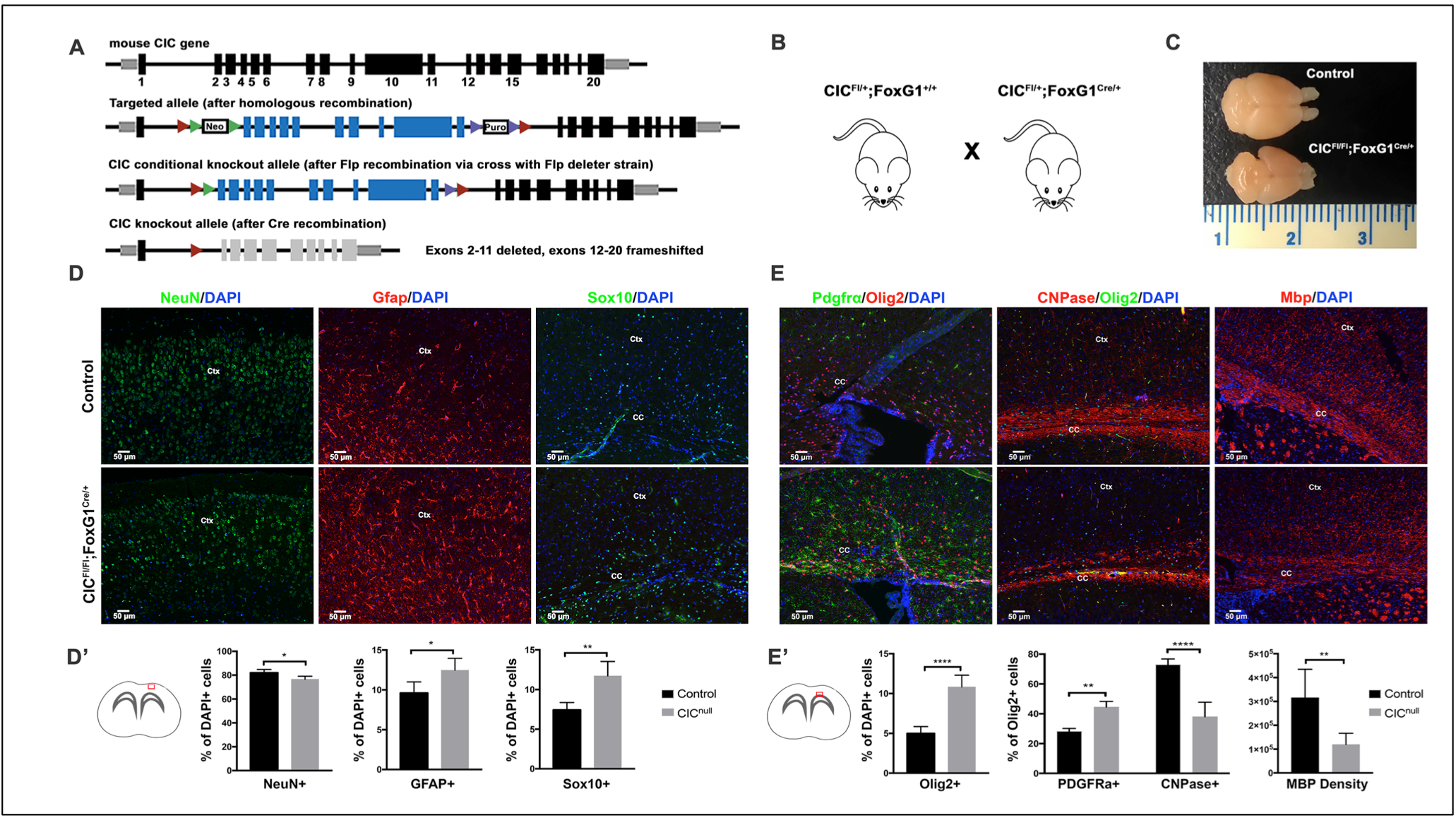
Forebrain-specific *Cic* deletion increases glial cells at the expense of neurons by P21. (A) Targeting strategy for Cic conditional knockout mice. Exon numbering is shown relative to Cic transcript variant 1. (B) Forebrain-deletion of Cic starting from E10.5 by crossing CIC-floxed line with FoxG1-cre. *Cic^Fl/Fl^;FoxG1^Cre^*^/+^ animals are compared with *Cic^Fl^*^/+^*;FoxG^Cre^*^/+^ or *Cic^Fl/Fl^;FoxG1*^+/+^ as controls. (C) Representative gross morphology of *Cic*-deleted and *Cic*-wildtype brains at P21. (D,D’) Staining and quantitation of NeuN+, Gfap+, and Sox10+ cells in Cic-deleted (*Cic^Fl/Fl^;FoxG1^Cre^*^/+^) cortex vs control (*Cic^Fl/Fl^;FoxG1*^+/+^). (E,E’) Staining and quantitation of Olig2+, Pdgfra+, and CNPase+ cells in lateral corpus callosum, and Mbp expression in lateral corpus callosum at P21. Results show quantitation from 5 mice per each group. Statistical analyses performed by unpaired t-test. Data shown as mean ± SD. *p<0.05, **p<0.01, ****p<0.0001. Ctx–cortex, CC–corpus callosum.

By crossing *Cic*-floxed mice to *Foxg1-Cre* mice (in which *Cre* is expressed throughout the forebrain by E10.5 (26)), we generated mice deficient in CIC in the embryonic telencephalon and subsequent postnatal cerebrum (Figure 2A,B). *Cic^Fl/Fl^;FoxG1^Cre^*^/+^ animals were born in approximate Mendelian ratios and were grossly indistinguishable at birth. They failed to thrive postnatally, however, becoming visible runts by postnatal day 7, and were uniformly lethal by P22 when they did not survive past weaning. The reason for lethality is unclear. Gross and cursory light microscopic exam showed the presence of the expected major anatomic structures in the brain, as well as the presence of laminated cortex, white matter tracts, deep nuclei, and hippocampi. The cerebra of Cic-null brains, however, were smaller than littermate controls (Figure 2C, Suppl Fig 2). We have not found major anatomic defects in other organs and suspect that poor feeding secondary to impaired neurologic function may be related to their progressive decline.

Microscopic exam showed that the overall decrease in cerebral size in CIC forebrain-null animals was due to global decreases in volume that affected both gray matter and white matter structures (Suppl Fig S2A-C). The corpus callosum of null animals was thinned but showed abnormally increased cellularity (Suppl Fig S2B). The cortex was also thinner in Cic-null brains, again showed increased cellularity (Suppl Fig S2C,D). Interestingly, cortical neuronal density was not significantly different between the two (Suppl Fig S2E), suggesting that the increased cellularity in the mutant brains could be due to increased glia, and also that the total number of cortical neurons was decreased (same neuronal density, but an overall smaller cortical volume).

Closer evaluation of P21 cortices confirmed this shift in cell populations, with decreased neurons and increased oligodendroglia cells and astrocytes in Cic-null cortices and white matter compared to controls (Fig 2D,D’,E,E’). In Cic-null cortices, NeuN+ cells comprised 76.6±2.5% of total cells; whereas in controls that were either CIC-floxed but without Cre (*Cic^Fl/Fl^; FoxG1*^+/+^) or that were heterozygous for Cic loss (*Cic^Fl^*^/+^*; FoxG1^Cre^*^/+^), NeuN+ cells comprised 82.04±2.78% of the total cells (n=5, p<0.05). In contrast, Gfap+ cells were increased in knockout relative to wild-type cortices (12.46±1.49 CIC-null *vs.* 9.61±1.39 Cic-wt; 1.29-fold change; n=5, p<0.05). Olig2+ cells were also increased in Cic-null cortices compared to controls (10.83±1.47% *vs.* 4.99±0.87%, in Cic-null vs control, respectively; 2.17-fold increased; n=5, p=0.0001), a finding corroborated by an increase in Sox10+ cells (11.72±1.83% *vs.* 7.43±0.94%; n=5, p=0.0001). In the progression along the oligodendrocyte lineage, the relative distribution of cells was also altered. Olig2 + Sox2+ cells were increased in Cic-null animals (CIC^Fl/Fl^;FoxG1^Cre/+^ 13.67±2.89% vs. Control 4.31±1.33%; n=3, p<0.01)(Suppl Fig S3) as were Olig2 + Pdgfra+ OPCs (44.53±3.80% of cells Olig2 + Pdgfra+ in CIC^Fl/Fl^;FoxG1^Cre/+^ *vs.* 28.20±1.95% in controls; p<0.01). In contrast, CNPase+ cells were comparatively decreased (*CIC^Fl/Fl^;FoxG1^Cre^*^/+^ 38.0±9.78% *vs.* Control 72.92±3.88%; n=5, p<0.0001) (Figure 2E,E’). Mbp expression in the white matter was also decreased in *Cic^Fl/Fl^;FoxG1^Cre^*^/+^ brains (n=6, p < 0.01) (Figure 2E,E’).

One possibility that could explain such a skewing of cell populations could be that there was increased death in a particular population. However we found no evidence of increased apoptosis in the knockout brains (Suppl Fig S4E). Other possibilities, were that Cic loss could be increasing proliferation and affecting lineage selection in NSCs, or that Cic loss could have specific effects on OPC proliferation and maturation – possibilities that are not mutually exclusive. In the following work, we focus our investigations on Cic’s role at the NSC stage, and examine its role in proliferation and lineage selection.

### CIC increases proliferation of neural stem cells

Others recently reported proliferative gains upon loss of Cic (13), and we too found evidence of this. By electroporating pCIG2-Cre (or pCIG2 empty vector control) into *Cic^Fl/Fl^* embryos, we could study the effects of Cic loss in a discrete population of VZ cells and their subsequent progeny (schematic Figure 4A). 48 hours after Cre electroporation into VZ NSCs at E13 (early to mid-neurogenesis), EdU incorporation was markedly increased, as were the numbers of Sox2 positive cells. Among GFP+ cells, EdU+ cells were increased by 2.6-fold in cre- vs. control-electroporated brains (pCIG2-Cre 9.99±0.92% Edu+, n=4, *vs.* pCIG2-Empty; 3.77±0.81%, n=5, p<0.0001)(Figure 3A,A’). The fraction of GFP+ cells expressing Sox2 was also increased (pCIG2-Cre 11.09±1.10% Sox2+ vs. pCIG2-Empty 5.48±1.01%; n=4 and n=5, respectively; p<0.05) (Figure 4B,B’). There was no change in activated Caspase-3, indicating that the increased Sox2+ fraction in Cic-deleted cells was not due to increased apoptosis in other cells (Suppl Figure S4D,D’). Together, these findings supported a cell-autonomous increase in NSC proliferation with CIC loss. Of note, there was also an increase in Edu+ cells among non-GFP cells within the area of the electroporated patches, suggesting additional non-cell autonomous effects that we did not pursue.

**FIGURE 3:**
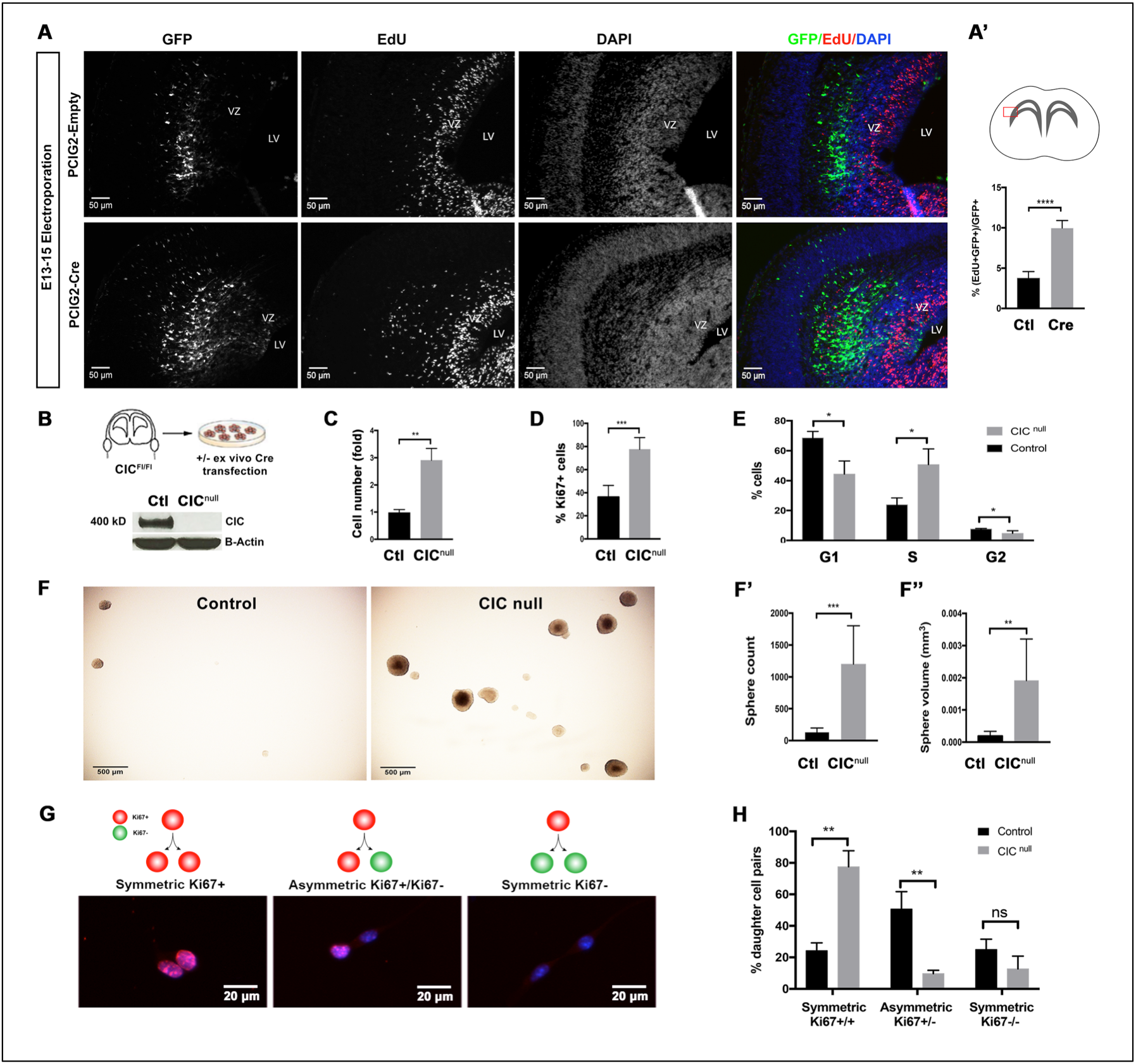
Cic deficiency increases proliferation and self-renewal of neural stem/progenitor cells. (A,A’) EdU incorporation 48-hours after electroporation of *Cre* or control plasmid into VZ of E13 *Cic-*floxed embryos; boxed area indicates zone for imaging and quantitation (n ≥ 4 mice per group). (B) Generation of Cic-null cells and control cells from *Cic*-floxed cells via ex vivo transfection of Cre recombinase, and western blotting for validation of knockout. (C) Trypan blue assay, (D) Ki67 proliferation index, and (E) propidium iodide cell cycle analysis in cultured cells. (F) Neural colony forming assay with quantitation of (F’) sphere number and (F’’) sphere size. (G) Paired cell assay schematic and representative images with Ki67 immunostaining and (H) Quantitation of symmetric proliferative, asymmetric, and symmetric terminal divisions. Data shown as mean ± SD. Data from ≥ 3 biologic replicates per condition. Statistical analysis performed by unpaired t-test. ns-not significant, *p<0.05, **p<0.01, ***p<0.001, ****p<0.0001. VZ–ventricular zone, LV–lateral ventricle.

**FIGURE 4:**
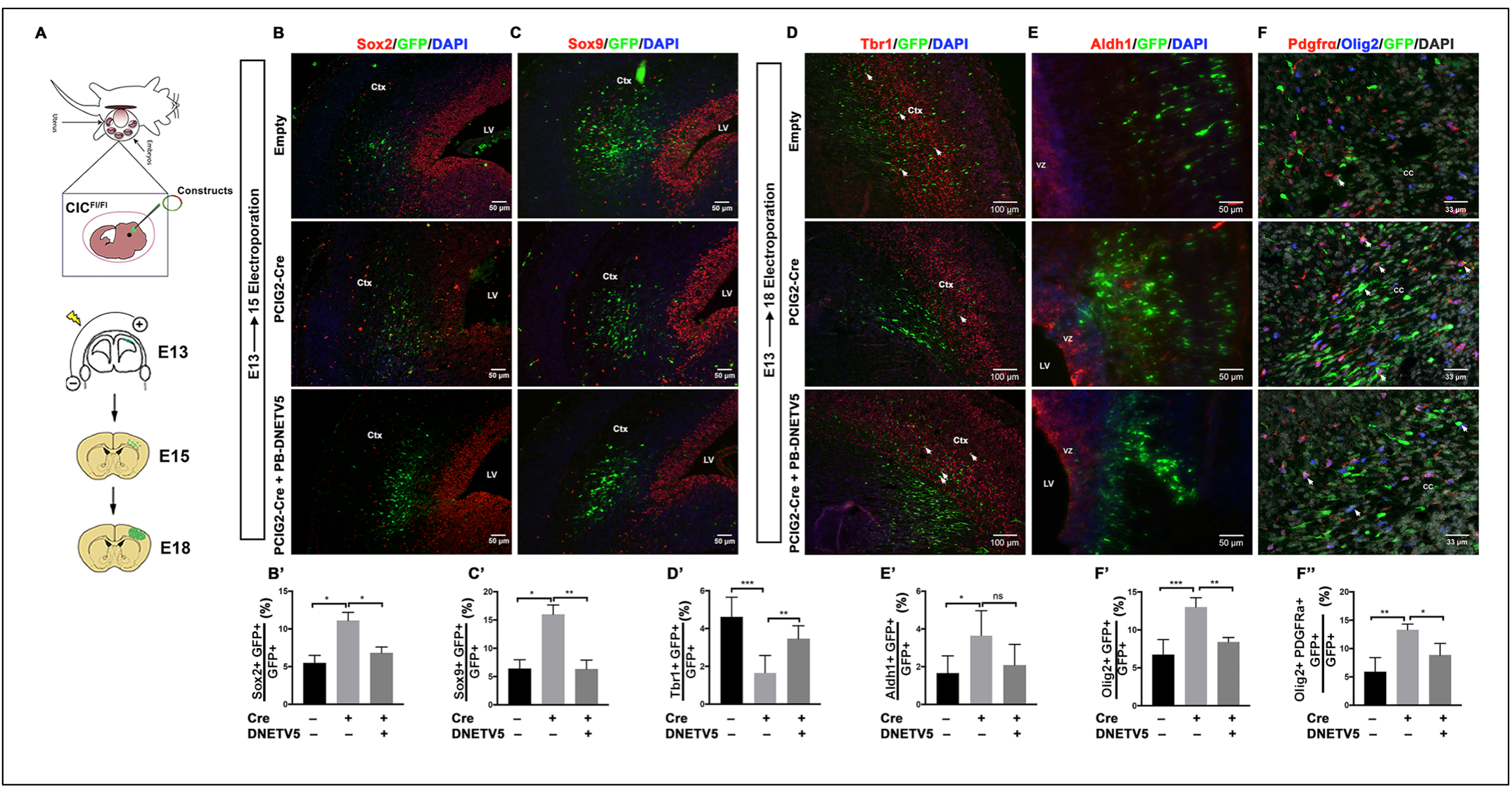
Deletion of *Cic* in the neurogenic period biases neural stem/progenitor cells to glial lineage selection. (A) Localized deletion of *Cic* in neural stem cells via *in-utero* electroporation into VZ cell of *C*ic-floxed embryos. Targeted cells carry a fluorescent marker and express either cre recombinase (Cre), dominant negative ETV5 (DNETV5), or not (Empty control), depending on the electroporated plasmid. Staining and quantitation of Sox2+ stem cells (B,B’) and Sox9+ glioblasts (C,C’) 2 days after E13 electroporation. Staining and quantitation of late-born Tbr1+ neurons, Aldh1+ astrocytes, Olig2+ oligodendrogial lineage cells, and Pdgfra+ oligodendrocyte precursors cells 5 days after E13 electroporation. Results show quantitation from 4 mice per group. Statistical analyses between control and experimental groups performed by ANOVA with Tukey’s posthoc test. Data shown as mean ± SD. *p<0.05, **p<0.01, ***p<0.001, ns–not significant. Ctx–cortex, LV–lateral ventricle, VZ–ventricular zone.

To confirm the cell autonomous gains in NSC proliferation with CIC ablation, we turned to cell culture. *Cic* null and control NSC lines were derived from E15 *Cic*-floxed cells transfected ex-vivo with Cre or control plasmids (Figure 3B). When maintained in serum-free NSC proliferation media, Cic-null and control cells both maintained expression of NSC markers such as the intermediate filament protein nestin (Figure 5A,B). When Cic-null and control cells were seeded at equal numbers and grown in NSC proliferation media, however, Cic-null cells displayed a 3-fold increase in cell numbers relative to controls by 3 days post-plating (Cic-null; 2.90±0.44, n=4, *vs.* Cic-wt; 0.98±0.12, n=3, p<0.01) (Figure 3C). These data were corroborated by Alamar blue cell viability assay (Suppl Figure S4A), Ki-67 immunostaining (Figure 3D) and cell cycle analysis (Figure 3E), all of which showed increased proliferation in Cic-null cultures compared to controls. There were no significant differences in the numbers of dead cells (Suppl Figure S4C), supporting that the differences in viable cell numbers for Cic-null cells was due to their higher proliferative rates, rather than reduced cell death. Together, the data show that Cic is a strong negative regulator of proliferation in forebrain NSCs.

**FIGURE 5:**
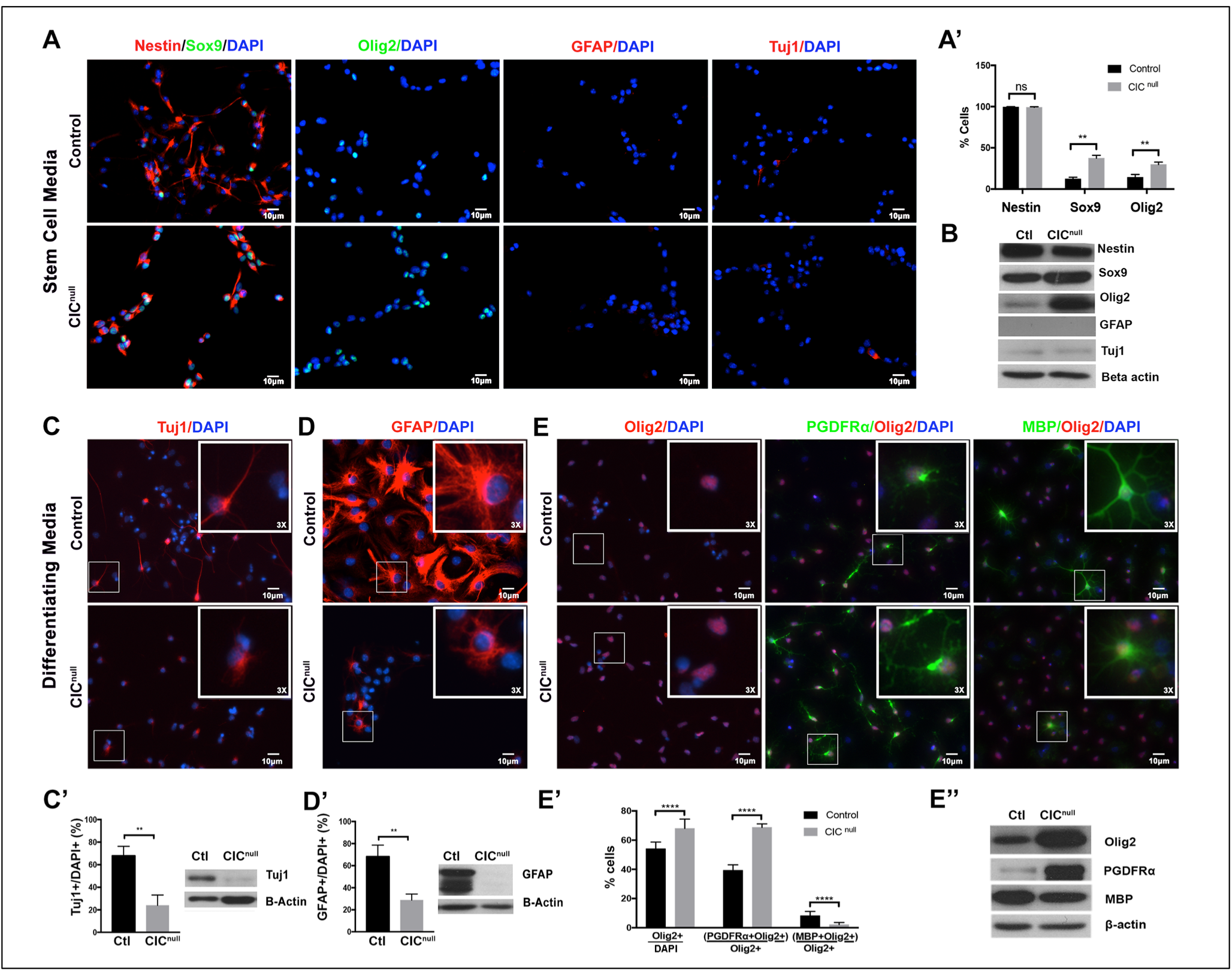
Cultured Cic null cells are oligodendrocyte lineage-biased. (A) Immunofluorescence and (B) Western blotting for Nestin, Sox9, Olig2, Gfap, and Tuj1 in Cic-null and control cells cultured in stem cell conditions. Responses of Cic-null and control cells to 10-day exposure to lineage-directed differentiation conditions for (C) neurons, (D) astrocytes, or (E) oligodendrocytes. Immunofluorescence quantitation of cell populations and western blotting of cells cultured in lineage-directed differentiation conditions for (C’) neurons, (D’) astrocytes, and (E’, E’’) stages of oligodendrocyte development. Data shown as mean ± SD. Data from ≥ 3 biologic replicates per condition. Statistical analysis performed by unpaired t-test. ns-not significant, **p<0.01, ****p<0.0001.

### *Cic* loss alters mode of cell division and self-renewal

During mammalian neurodevelopment, the stem cell pool is first expanded by symmetric proliferative cell divisions followed by rounds of asymmetric divisions of stem and progenitor cells, which generate neurons and then glia, and finally by terminal symmetric differentiative divisions (18). The balance between symmetric and asymmetric divisions is important for maintaining the stem/progenitor pool, and alterations in cell division mode can lead to neoplasia (27).

To investigate whether *Cic* loss alters modes of cell division, we performed a paired cell assay. *Cic* null and control NSCs were seeded at low density in adherent cultures for 24 hrs, then fixed and stained for Ki-67 (Figure 3G). Pairs where both daughter cells were Ki67+ were scored as symmetric proliferative divisions. Pairs where daughter cells differed in Ki67 expression (i.e. Ki67+/Ki67−) were scored as asymmetric. Pairs where both daughter cells were Ki67− were scored as symmetric terminal. *Cic* null cells underwent significantly more frequent symmetric proliferative divisions (77.6 ± 10.2% vs. 24.3 ± 4.9%; p<0.01, n=3) and fewer asymmetric divisions (9.7 ± 1.2% vs. 64.0 ± 7.1%; p < 0.05, n=3) compared to controls (Figure 3H). No significant difference was found in numbers of symmetric terminal divisions between *Cic* null and control cells, although there was a trend toward decreased terminal divisions in CIC null cells (12.7 ± 8.1% vs. 25.0 ± 6.5%; n=3, p = 0.108)(Figure 3H). The net effect of these is the presence of more cycling and self-renewing cells.

Consistent with this, evidence for increased self-renewal in the Cic null cells was also detected in a clonogenic assay (Figure 3F). In this assay, after plating equal numbers of dissociated cells in semi-solid media, a higher number of spheres indicates higher number of self-renewing cells in the initial population, whereas sphere volume is a more general indicator of proliferation that includes the influence of factors such as cell cycle kinetics, modes of cell division, and the fraction of cells remaining in or exiting the cell cycle. Both Cic-null and control NSCs were able to generate spheres, but Cic-null NSCs NSCs generated higher sphere counts (n=3; p < 0.0001) and larger spheres compared to control (n=3; p<0.0001)(Figure 3F,F’,F’’). Thus, Cic loss confers not only higher proliferation but higher self-renewal capacity in NSCs, at least when cells are in an environment promoting NSC proliferation.

### *Cic* ablation in NSCs increases the pool of cells expressing glial lineage determinants

While the proliferative effects of Cic deficiency in NSCs were clear, what was intriguing was the additional possibility that Cic loss may be affecting lineage selection in NSCs. As the *Cic^Fl/Fl^;FoxG1^Cre^*^/+^ phenotype was characterized by a shift in cell populations with fewer neurons and increased glia (Figure 2), we thus asked whether Cic loss had altered the expression of factors important in regulating lineage selection and gliogenic potential.

To this end, we examined the expression of Sox 9 and Olig2 in Cic-wildtype and -null NSCs. Sox9 is an HMG-box transcription factor present in a range of CNS cell types, including stem cells, astrocyte and oligodendroglial precursors, and later glial cells. It has key roles not only in stem cell maintenance but also in driving differentiation programs away from neurogenesis and towards gliogenesis of both astrocytes and oligodendrocytes at the stage of gliogenic initiation (28–30). Similarly, Olig2 is expressed in a multitude of cells including stem cells, oligodendroglial lineage cells, and specific subtypes of neurons; but it has a major role in establishing oligodendroglial competence. In our control or cre-electroporated brains where Cre was introduced at E13, we found that the fraction of GFP+ cells expressing Sox9 was 2.5-fold increased over controls at 2 days post-electroporation (pCIG2-Cre; 15.97±1.69% *vs.* pCIG2-Empty; 6.37±1.60%, n=4 p<0.05) (Figure 4C,C’). Following the targeted cells to E18 (5 days post-electroporation) revealed that a smaller fraction Cic-deleted NSCs subsequently became Tbr1+ early-born/deep-layer neurons (pCIG2-Cre 1.66±0.91% *vs.* pCIG2-Empty 4.62±1.04%; n=5, p<0.01) (Figure 4D,D’). Conversely, a greater number fraction became Aldh1+ astrocytes (pCIG2-Cre 3.65±1.31%, n=5, *vs.* p-CIG2-Empty 1.664±0.91%, n=6; p<0.05) or Pdgfra+ OPCs (pCIG2-Cre 13.29±1.04%, *vs.* p-CIG2-Empty 5.87±2.54%; n=4; p<0.01) (Figure 4C,C’,E,E’).

These findings were echoed in the expression of stem and lineage markers in the cultured NSCs. As expected, when cells were grown in NSC media, both Cic-null and control cells both strongly expressed Nestin, and were largely devoid of Gfap or the pan-neuronal marker βIII-Tubulin (as detected by Tuj1)(Figure 5A,B), or markers of subsequent oligodendrocyte stages, including OPCs, mature, and myelinating oligodendrocytes (not shown). There, was, however, a marked increase in the percentage of cells expressing Sox9 and Olig2 in Cic-null cultures (Figure 5A,A’). Sox9+ cells were increased 3.04-fold in Cic-null cultures compared to control (Cic-null 37.4±3.6% *vs.* Cic-wt 12.3±2.1%; n=3, p < 0.01)(Figure 5A,B) while Olig2+ cells were increased 2.09-fold (Cic-null 29.8±2.9% *vs.* Cic-wt 14.2±3.5% in controls; n=3, p<0.01) (Figure 5A,A’). Western blotting also corroborated the findings (Figure 5B). A possible interpretation of the increased Sox9 and Olig2 concurrent with Nestin positivity in Cic-null cells is that Cic loss may set an intrinsic foundation for pro-glial or pro-oligodendroglial programs starting early in the neural cellular hierarchy.

### Cic-deficient NSCs are less responsive to extrinsic neuronal and astrocytic lineage selection cues, but are biased to oligodendroglial lineage selection

To directly test cell type specification capacity, we challenged Cic-null and control NSCs with exposure to different lineage-promoting culture conditions. Neuronal differentiation was induced by culturing cells with B27 and cAMP. Astrocytic differentiation was induced by culturing NSCs in 1% FBS and N2. Oligodendroglial differentiation was induced by culturing cells in media with B27 and tri-iodo-thyronine. After a 10 day exposure to these conditions, cultures were analyzed for cellular identity and morphology.

After 10 days in the neuronal condition, fewer Tuj1+ cells were generated in absolute numbers in Cic null cultures compared to controls after plating equal numbers of starting cells (1.967±0.36 x10^3^ vs 2.929±0.33 x10^3^ Tuj1+ per well in Cic-null vs control; p<0.05)(Fig 6G) – a finding corroborated by western blotting for Tuj1 (Figure 5C’). Furthermore, of the cells that were Tuj1+, we observed fewer and less complex cell processes in the Cic-null cells than in their Cic-wildtype counterparts (Figure 5C). In the astrocytic condition, there were fewer absolute numbers of Gfap+ cells in the Cic-null cultures (1.471±0.15 x10^3^ vs 2.74±0.55 x10^3^ Gfap+ per well in Cic-null vs control, p<0.05) (Fig 6F). Western blotting also showed decreased GFAP (Figure 5D’). Analogous to our observation in the neuronal differentiation condition, Gfap+ Cic-null cells in the astrocytic differentiation condition displayed more rudimentary processes than their Cic-wildtype counterparts (Figure 5D). In both of these conditions, there were only a few proliferating cells remaining after the differentiation protocols, consistent with successful differentiation. In both conditions, there was also no change in apoptotic cells (Suppl Figure S4F-F’’’). When viewed as fractions of the total cell population, Cic-null cultures were comprised of a significantly smaller fraction of bIII-Tub+ (Tuj1+) neurons in the neuronal condition (CIC-null 24.0±9.2% *vs.* CIC-wt 67.8±8.5%, respectively; n=3, p<0.01) and a significantly smaller fraction of Gfap+ cells in the astrocytic condition (29.1±5.5% in Cic-null *vs.* 68.2±10.4% in Cic-wt; n=3, p <0.01)(Figure 5C,D). Thus, with normal CIC expression, NSCs that are exposed to neuronal conditions become neurons (as expected), but CIC-deficient NSCs do not as readily select the neuronal lineage when exposed to the same neuronal-inducing conditions. Likewise, most CIC-wt NSCs that are exposed to astrocytic conditions become astrocytes (as expected), but CIC-null NSCs do not as readily select the astrocytic lineage when exposed to the same astrocyte-inducing conditions. Together, the data indicate that Cic-deficient NSCs are less responsive to external cues for neuronal and astrocytic lineage specification.

**Figure 6:**
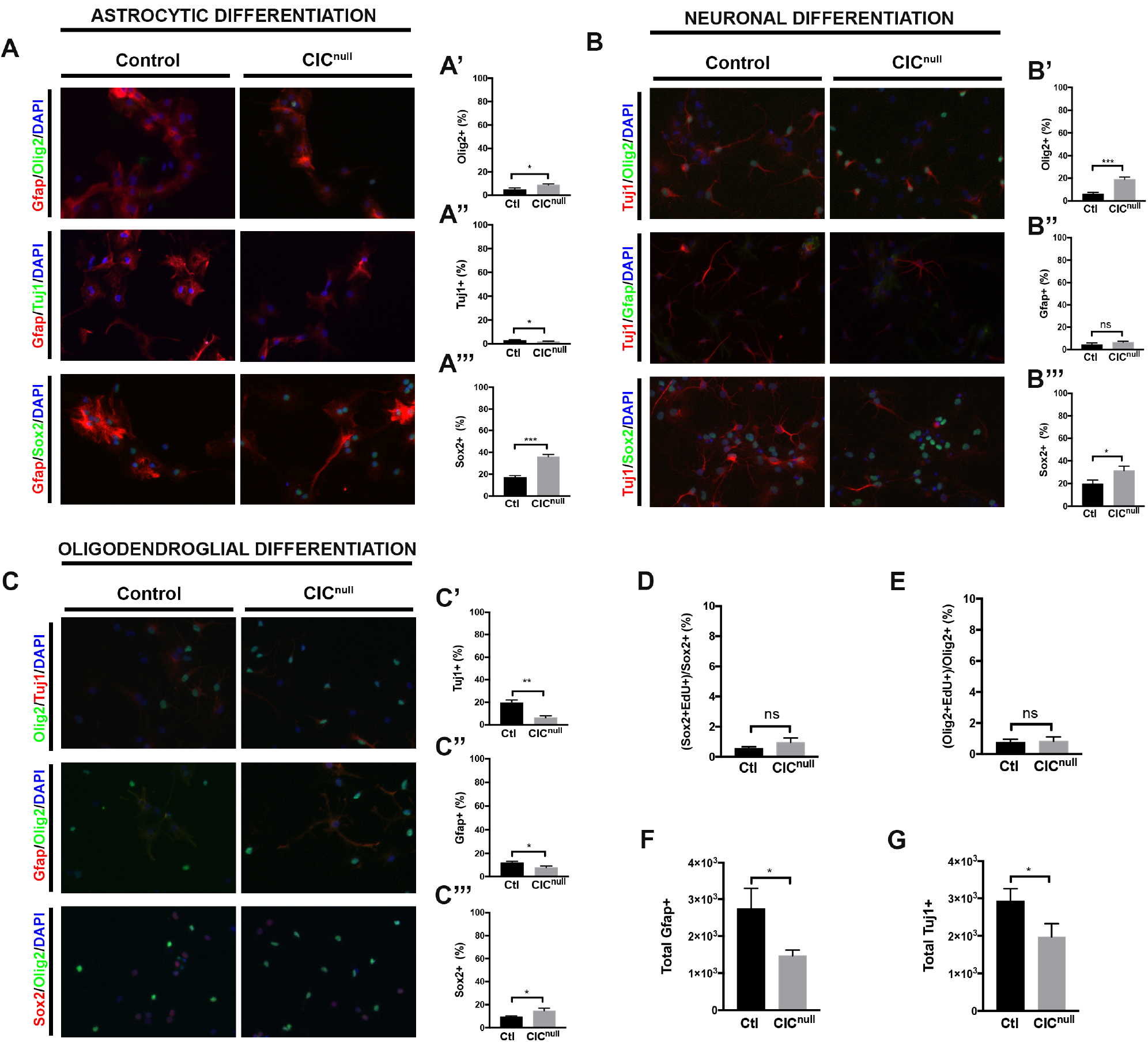
Alternate lineage selection of Cic null NSCs in response to extrinsic differentiation cues. (A) Under astrocytic promoting conditions, representative images of cultures stained for oligodendrocytes (Olig2+), neurons (Tuj1+) and stem cells (Sox2+); and (A’,A’’,A’’’) corresponding quantifications. (B) Under neuronal promoting conditions, representative images of cultures stained for oligodendrocytes (Olig2+), astrocytes (Gfap+) and stem cells (Sox2+); and (B’,B’’,B’’’) corresponding quantifications.(C) Under oligodendrocytic promoting conditions, representative images of cultures stained for neurons (Tuj1+), astrocytes (Gfap+) and stem cells (Sox2+); and (C’,C’’,C’’’) corresponding quantifications. (D) Percentage of Sox2+ cells positive for Edu incorporation after 10 days exposure to neuronal promoting conditions. (E) Percentage of Olig2+ positive for Edu incorporation after 10 days exposure to neuronal promoting conditions. (F) Absolute counts of Gfap+ cells/well after differentiating 4×10^3^ NSCs in astrocytic conditions for 10 days. (G) Absolute counts of Tuj1+ cells/well after differentiation 4×10^3^ NSCs in neuronal conditions for 10 days. Bars indicate mean ± SD. ADC-Astrocytic differentiation condition, NDC-Neuronal differentiation condition, ODC-Oligodendroglial differentiation condition. ns – not significant, * p<0.05, ***p<0.001.

We then asked what the Cic-null NSCs became, if not neurons and astrocytes, respectively. In the neuronal condition, the Tuj1-population was predominantly comprised of Sox2+ cells, followed by Olig2+ cells; together, half of the culture was comprised of either Sox2+ (31%) or Olig2+ cells (19%) after exposure to the neuronal conditions (Figure 6B’,B’’’). Of the Sox2+ and Olig2+ cells present, however, less than 1% were EdU+, and there was no significant difference between the proliferation rate of Sox2+ or Olig2+ cells in the Cic-null vs. control cultures in the neuronal condition (Figure 6D-E). There was also a small but non-significant increase in the percentage of Gfap+ cells in the Cic-null cultures compared to control. In the astrocytic culture condition, the Gfap-population was also comprised predominantly of Sox2+ cells and, to a lesser extent Olig2+ cells, both of which were significantly increased in the Cic-null cultures compared to controls. 36% of the Cic-null cells in the astrocytic condition still expressed Sox2 after the 10-day protocol (Figure 6A’’’). Tuj1+ cells were rare in the astrocytic condition in both Cic-null and control cultures. These data further support that CIC-deficient NSCs are less responsive to neuronal and astrocytic lineage cues, and instead remain as stem cells that are permissive to the oligodendroglial lineage.

After 10 days in the oligodendrocyte-promoting media both Cic-null and wildtype cultures contained a mix neurons, astrocytes, and oligodendrocytes; however the distributions of cells between cell types and also along stages of oligodendrocyte lineage progression were altered. There was a higher fraction of Olig2+ cells in Cic-null cultures compared to control (Cic-null 68.0±1.6% *vs.* Cic-wt 54.1±0.9%; n=3, p < 0.01) as well as increased Olig2 by western blotting (Figure 5E). When analyzed for markers of OPCs versus more mature oligodendroglial markers, Cic-null cells also had a greater percentage of Olig2+ Pdgfra+ cells than in Cic-wildtype cultures (Cic-null 68.8±2.4% vs. Cic-wt 39.3±3.8%; n=5, p < 0.0001)(Figure 5E). Conversely the percentage of Olig2+MBP+ cells was decreased in Cic-null cultures compared to controls (Cic-null 2.1±1.5% vs. Cic-wt 8.2±3.0%; n=5, p < 0.0001)(Figure 5E). Of the few Cic-null Olig2+MBP+ cells that were identified, process formation was rudimentary compared to Cic-wildtype cells. With respect to other cell types present in the oligodendroglial culture conditions, Sox2+ cells were increased in Cic-null cultures compared to control, whereas both Tuj1+ and Gfap+ cells were reduced (Figure 6C-C’’’).

The results suggest to us that Cic-null cells are less sensitive to neuronal or astrocytic differentiation signals but retain permissiveness to the oligodendroglial lineage. Cumulatively, based on the pattern of cell-type specific nuclear Cic expression in forebrain tissue together with the in vivo and in vitro functional studies of Cic ablation, we conclude that Cic is a critical regulator of proliferation, self-renewal, and cell fate of NSCs. Loss of Cic expands the OPC pool by not only increasing NSC proliferation but also biasing their specification towards the oligodendroglial lineage.

### Ets factors are transcriptional repressive targets of Cic in the forebrain

As Cic is a transcriptional repressor, one mechanism for our findings is that Cic loss de-represses specific genes driving NSC proliferation and lineage specification. In this respect, *Etv* genes, which encode Ets-domain transcription factors, are candidate target genes of interest. *Etv* genes (Etv 1, 4, 5) have been identified as direct targets of Cic in various mammalian cells and tissues (12, 31), and are overexpressed in ODGs (32). Furthermore, previous studies have documented Cic occupancy at the *Etv5* promoter, at least in cerebellar tissue (33). Consistent with a possible functional relationship between CIC and Ets factor gene regulation in the forebrain, we found that NSCs and OPCs, the two cell types that have the lowest levels of nuclear Cic, express the highest levels of Etv4 and 5 (Figure 7F-K; Supplemental Figure S5B-D’). We also confirmed that in forebrain tissue, *Etv5* showed evidence of promoter occupancy by Cic via ChIP-PCR (Figure 7C).

**FIGURE 7:**
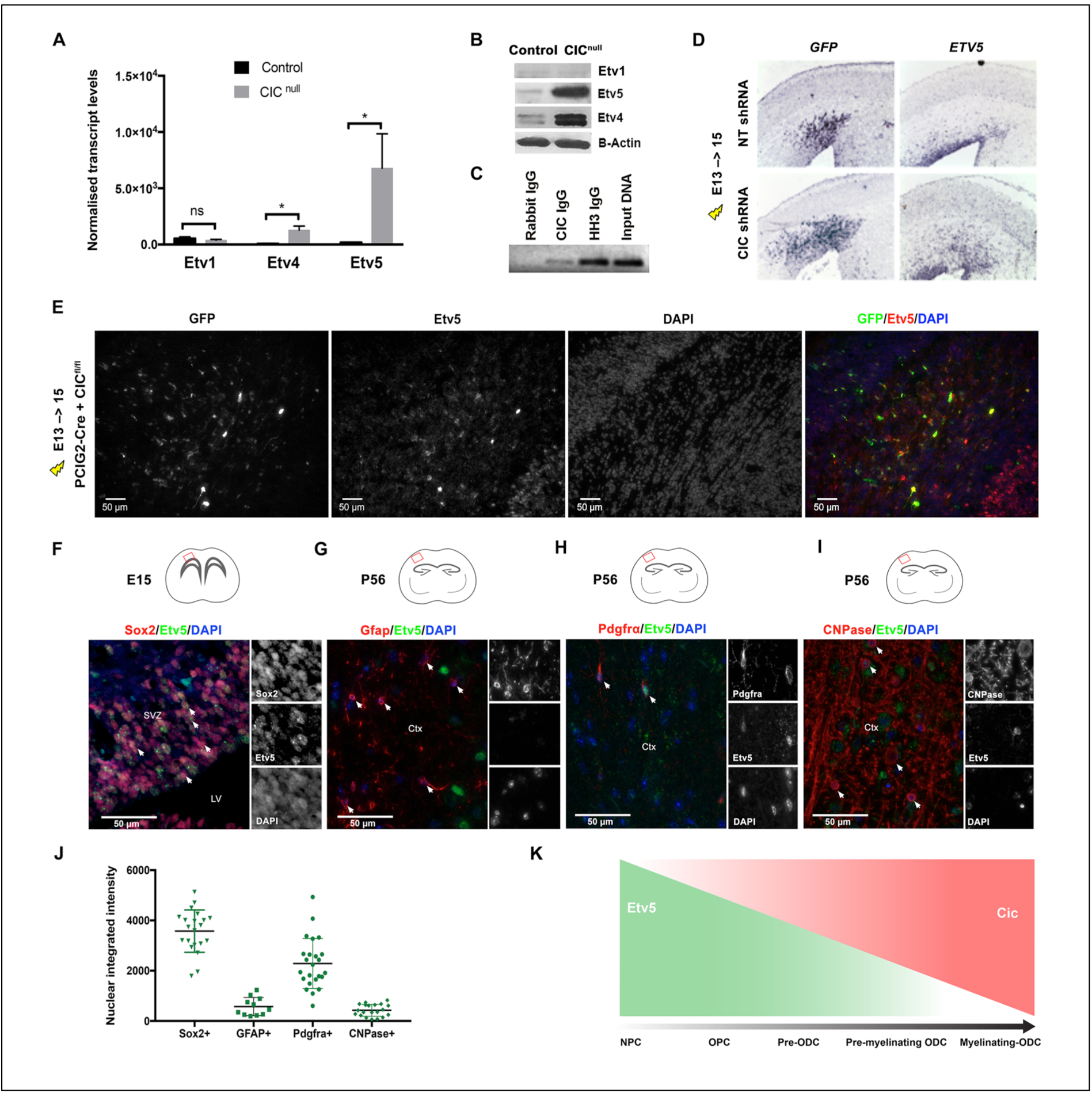
*Etv5* is a direct target of Cic transcriptional repression. *Etv1*, *Etv4*, and *Etv5* mRNA (A) and protein (B) levels in cultured CIC-null and -control cells after 48-hour exposure to oligodendrocytic differentiating conditions; transcript levels normalized to average of 3 (Actin, GAPDH, Tubulin beta chain) housekeeping genes. Data from n=3 biologic replicates. (C) ChIP-PCR for Cic at the *Etv5* promoter. (D) Electroporation of Cic shRNA or non-targeting (NT) shRNA at E13 followed by *in-situ* hybridization for *Etv5* and *GFP* at E15; *Etv5* transcripts are upregulated in areas of Cic knockdown. Data from n ≥ 3 mice per group. (E) Immunofluorescence staining showing Etv5 and GFP protein expression 2 days after Cre electroporation into E13 VZ of *Cic*-floxed embryos. *Cic* deleted cells show increased Etv5 protein. (F-I, J) Differential Etv5 expression in (F) NSCs (Sox2+) at mid-neurogenesis, and in (G) astrocytes (Gfap+), (H) OPCs (Pdgfra+), and (I) mature oligodendrocytes (CNPase+) in P56 adult cortex. Each point represents one cell quantitated. Results show quantitation from ≥20 cells for each marker. Data shown as mean ± SD. Statistical analyses performed by unpaired t-test. *p<0.05. SVZ–subventricular zone, LV–lateral ventricle, Ctx–cortex. (K) Schematic depiction of relationship of Cic and Etv5 expression as neural stem cells differentiate to mature myelinating oligodendrocytes.

If Ets factors are the mediators of the effects of CIC loss that we observe, we would expect their levels to be elevated in our experimental systems of CIC loss. Indeed, we found that *Etv4* and *Etv5* transcript levels were elevated in cultured Cic-null cells compared to control cells after 2 days of growth in oligodendroglial conditions. *Etv5* mRNA was increased ∼37 fold while *Etv4* transcripts were increased ∼19 fold in Cic-null cells versus control cells whereas *Etv1* transcripts were unchanged (Figure 7A). Western blotting also confirmed marked increases in levels of Etv4 and Etv5 in Cic-null cells (Figure 7B). Although both *Etv4* and *Etv5* were both de-repressed upon Cic loss, because of the comparatively higher levels of Etv5 relative to Etv4, as well as previous studies implicating *Etv5* in mediating glial fate decisions (34), we focused additional studies on this Ets factor. In vivo electroporation of *Cic* shRNA into the telencephalic VZ resulted in upregulation of *Etv5* transcript in the electroporated patch within 48 hours (Figure 7D). Our *Cic*-floxed, Cre-electroporated brains showed a similar increase in Etv5 protein in the electroporated patch (Figure 7E). In aggregate, these data support that the PEA3 Ets transcription factors are transcriptional repressive targets of Cic in the forebrain.

### Ets de-repression mediates the proliferative and oligo-biased phenotype of *Cic*-deficient NSCs

To determine whether the increased proliferation and OPC bias that we observed with CIC ablation was mediated by Etv5, we overexpressed wild-type *Etv5*. In vivo, electroporation of wild-type *Etv5* phenocopied the increase proliferation of CIC-null cells in vivo (Figure 8D, D’, S6A). Similarly, *Etv5* overexpression acutely increased proliferation of cultured NSCs (Figure 8F, S6C’). We also performed epistasis experiments introducing a dominant negative form (*DNETV5*) in which *Etv5* was fused to the *Engrailed* transcriptional repressor domain. Introduction of *DNETV5* or knockdown of *Etv5* with siRNA reduced the proliferation of cultured *Cic-*deficient NSCs back to control levels (Figure 8F-G, Suppl Figure S6D). In vivo, the increased proliferation observed upon deletion of *Cic* by cre-electroporation was abrogated by co-electroporation with *DNETV5,* resulting in proliferation that was comparable to baseline (Figure 8E, E’ compared to 8D,D’; Figure S6). Based on the extent of the effects of *Etv5* overexpression and *DNETV5* rescue in these assays, we conclude that the proliferative effects of Cic loss are large driven by de-repression of Etv5.

**FIGURE 8:**
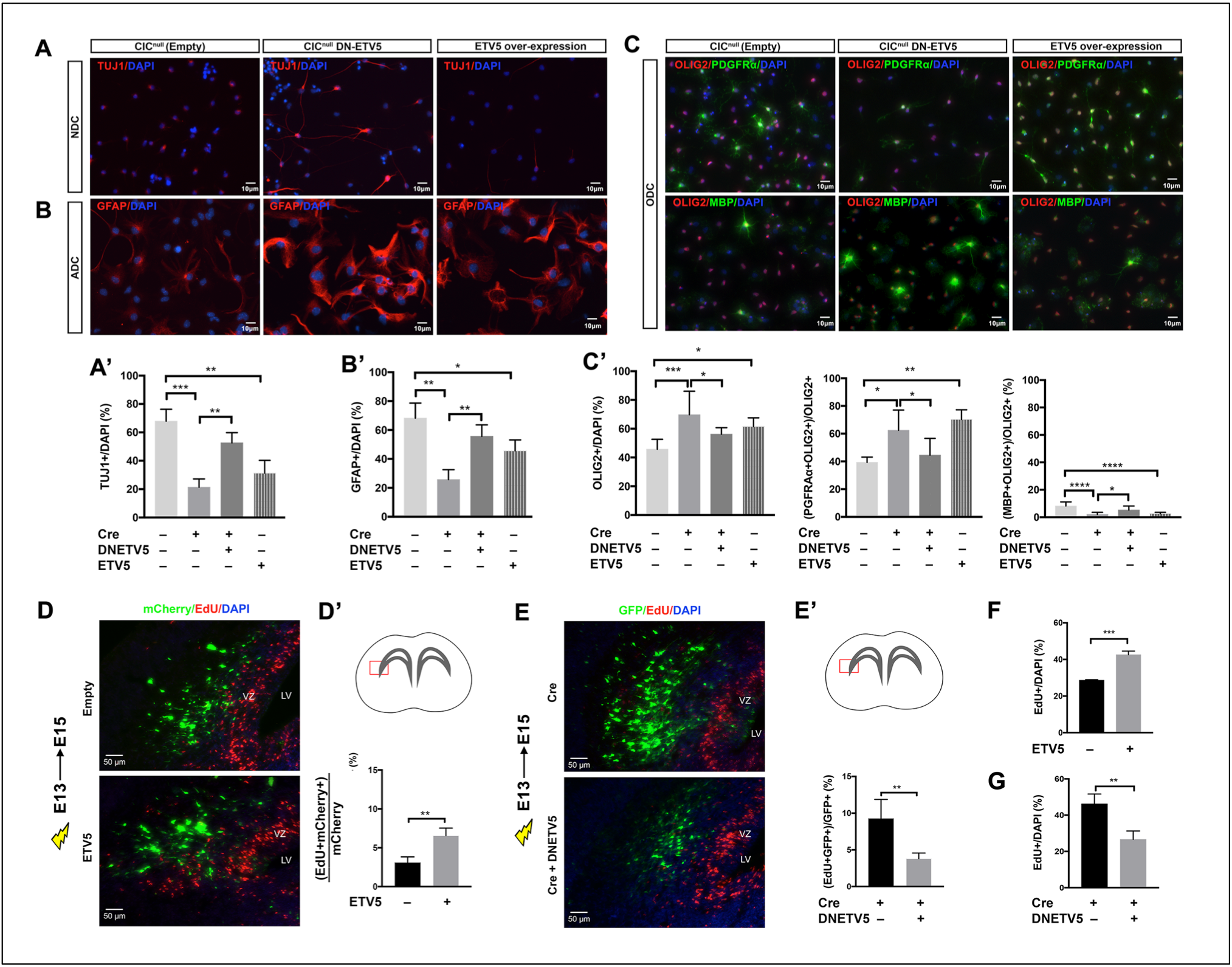
Etv5 is necessary and sufficient for proliferation and cell fate bias effects downstream of Cic loss. Cic-null cells with and without DN-ETV5, and Cic-wildtype cells with Etv5 overexpression were assessed for their ability to differentiate in response to 10-days’ exposure to lineage-directed differentiating conditions. (A,A’) Neuronal differentiation capacity was assessed by bIII-Tubulin immunostaining (Tuj1 positivity), (B,B’) astrocytic differentiation capacity was assessed by Gfap immunostaining, and (C,C’) oligodendroglial differentiation capacity was assess by staining for Olig2, Pdgfra, and Mbp. (D,D’) EdU incorporation 2 days post-electroporation of wildtype *ETV5* or empty control plasmid, both carrying mCherry as a marker, into E13 CIC^Fl/Fl^ VZ. Note: mCherry fluorescence and EdU staining were false-colorized to green and red after grayscale imaging. (E,E’) EdU incorporation 2 days post-electroporation of Cre only or of Cre co-electroporated with *DN-ETV5* into E13 CIC^Fl/Fl^ VZ. EdU incorporation assay in cultured NSCs showing (F) effects of ETV5 overexpression in CIC-wildtype NSCs and (G) effects of *DN-ETV5* expression in Cic-null NSCs. Data shown as mean ± SD from n ≥ 4 mice per each group for in vivo studies and n=3 biological replicates for cell culture studies. Statistical analyses performed either t-test in D’, E’, F, G; or with ANOVA with Tukey’s posthoc test in A’, B’, C’. ns-not significant, *p<0.05, **p<0.01, ***p<0.0001. ADC–astrocytic differentiation condition, NDC–neuronal differentiation condition, ODC–oligodendrocytic differentiation condition. VZ–ventricular zone, LV–lateral ventricle.

With respect to the altered proportions of cell types that we had observed in CIC-deficient NSCs exposed to the different lineage-specific culture conditions, overexpression of wild-type Etv5 also phenocopied CIC loss in many respects. There were decreased fractions of Tuj1+ cells and Gfap+ cells in the neuronal and astrocytic conditions, and increased fractions of Olig2+ and Olig2+Pdgfra+ cells in the oligodendrocytic condition when Etv5 was overexpressed – although the severity of the changes was somewhat less marked with Etv5 overexpression compared to Cic loss. DNETV5 was also able to substantially (although incompletely) rescue the phenotype of Cic-deficient cells in these assays. In the neuronal and astrocytic conditions, DNETV5 significantly increased the populations of Tuj1+ and Gfap+ cells, respectively. The shifts in Olig2+ cells and Olig2+Pdgfra+ OPCs returned to control levels (44.6±12.0% in *Cic^null^;DNETV5* cells *vs.* 62.5±14.6% in *Cic^null^* cells; n=3, p<0.05). The percentage of cells expressing MBP also showed a substantial although partial rescue (5.3±2.9% in *Cic^null^;DNETV5* cells *vs.* 2.1±1.5% in *Cic^null^* cells; n=3, p < 0.05) (Figure 8C-C’). That the extent of the changes for the lineage selection assays with Etv5 overexpression and DNETV5 rescue were partial suggests that other factors may contribute to the lineage specification affects, nevertheless the results point to Etv5 as playing a large part in the lineage phenotype.

Thus, Etv5 is both necessary and sufficient for inducing the proliferation and cell fate bias effects downstream of CIC loss.

### Ets blockade decreases proliferation, self-renewal, and tumorigenicity of human oligodendroglioma cells

Finally, we used 2 oligodendroglioma cell lines, BT54 and BT88, to further investigate the importance of *CIC* mutation/loss in human disease. Both of these patient-derived lines harbour the characteristic whole-arm chromosome 1p and 19q losses that are diagnostic of ODG (35). As well, BT54 harbours a splice acceptor site mutation in the remaining *CIC* exon 6 while BT88 harbours a missense mutation in the remaining *CIC* exon 20 (3). In both BT54 and BT88, either stable re-expression of wildtype CIC or stable expression of *DNETV5* significantly reduced proliferation as measured by EdU incorporation (Figure 9A,A’,C). Sphere-forming ability was also decreased by *CIC* expression or by introduction of *DNETV5* (Figure 9B,B’,B’’,D,E). These results were also confirmed by transient siRNA experiments to knock down Etv5 in BT-88 cells that also showed decreased proliferation and sphere formation (data not shown). Consistent with an ongoing requirement for CIC loss or Etv5 expression for proliferation and self-renewal in ODG, in vivo tumorigenicity of BT88 cells was markedly reduced, and survival of animals was increased, by stable expression of either wildtype *CIC* or *DNETV5* (Figure 10). Thus, in cells that are already transformed to ODG, CIC loss and the subsequent elevation of Etv5 remains important to sustain the proliferative phenotype.

**FIGURE 9:**
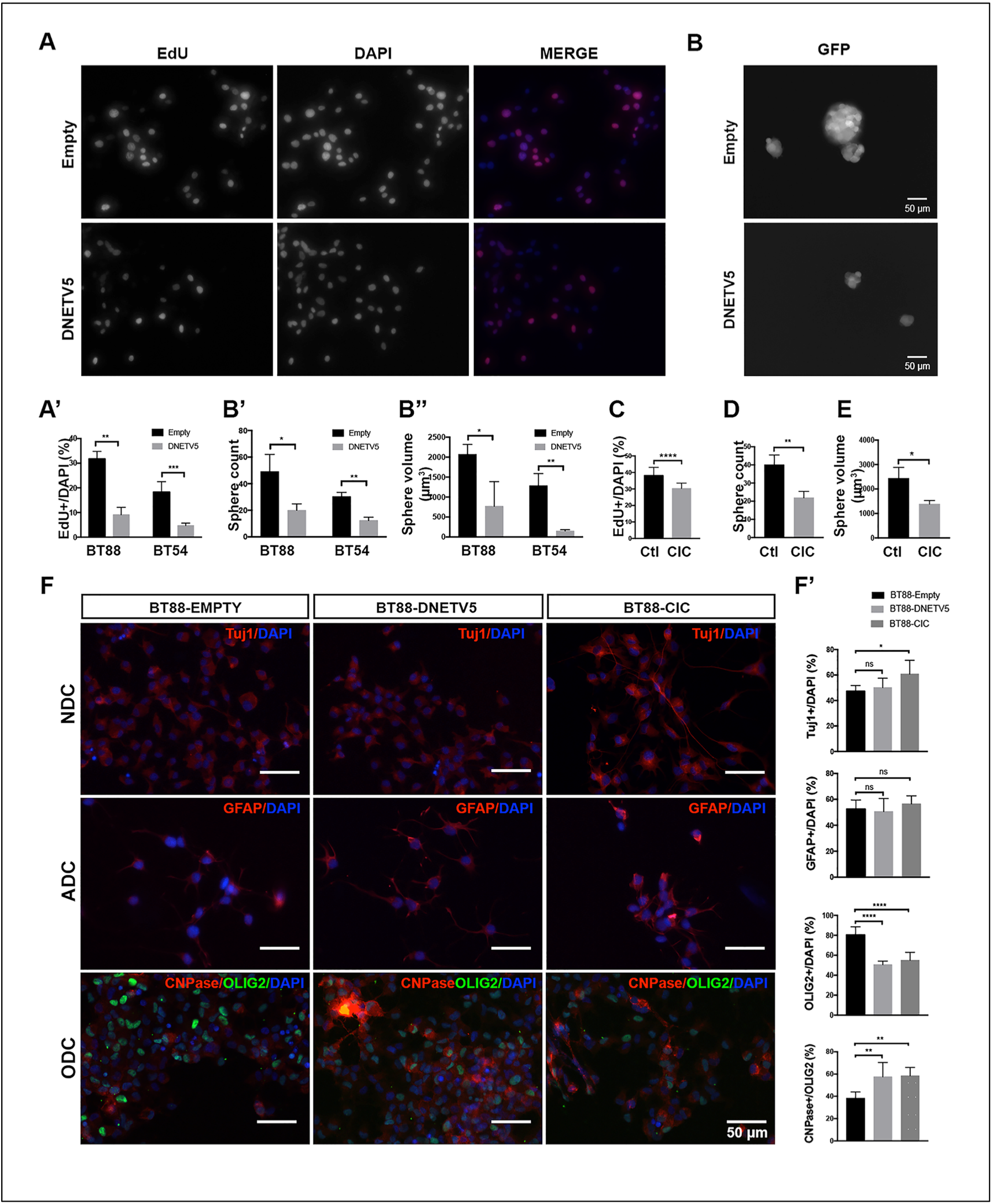
ODG cells require CIC loss or elevated ETV5 to maintain proliferation and stemness. (A,A’) EdU incorporation in BT-88 and BT-54 ODG cell lines stably transfected with control plasmid or of DNETV5. Quantitation shows results from both cell lines. Representative images are from BT-88. (B) Representative images showing effect of DN-ETV5 on sphere formation in BT88 ODG cells. Quantification of (B’) sphere number and (B”) size in BT-54 and BT-88 ODG cells without and with DNETV5. (C,D,E) CIC re-expression in BT-88 cells decreases proliferation, sphere number, and sphere size. (F,F’) Lineage-specific differentiation capacity of BT-88 ODG cells stably transfected with either control plasmid, DNETV5, or CIC; expression of Tuj1, GFAP, OLIG2, and CNPase after 10-days exposure to differentiation conditions for neurons, astrocytes, and oligodendrocytes. Data shown as mean ± SD from n=3 biologic replicates. Statistical analyses by ANOVA with Tukey’s posthoc test. ns-not significant, *p<0.05, **p<0.01, ****p<0.0001. NDC–neuronal differentiation condition, ADC–astrocytic differentiation condition, ODC–oligodendrocytic differentiation condition.

**FIGURE 10:**
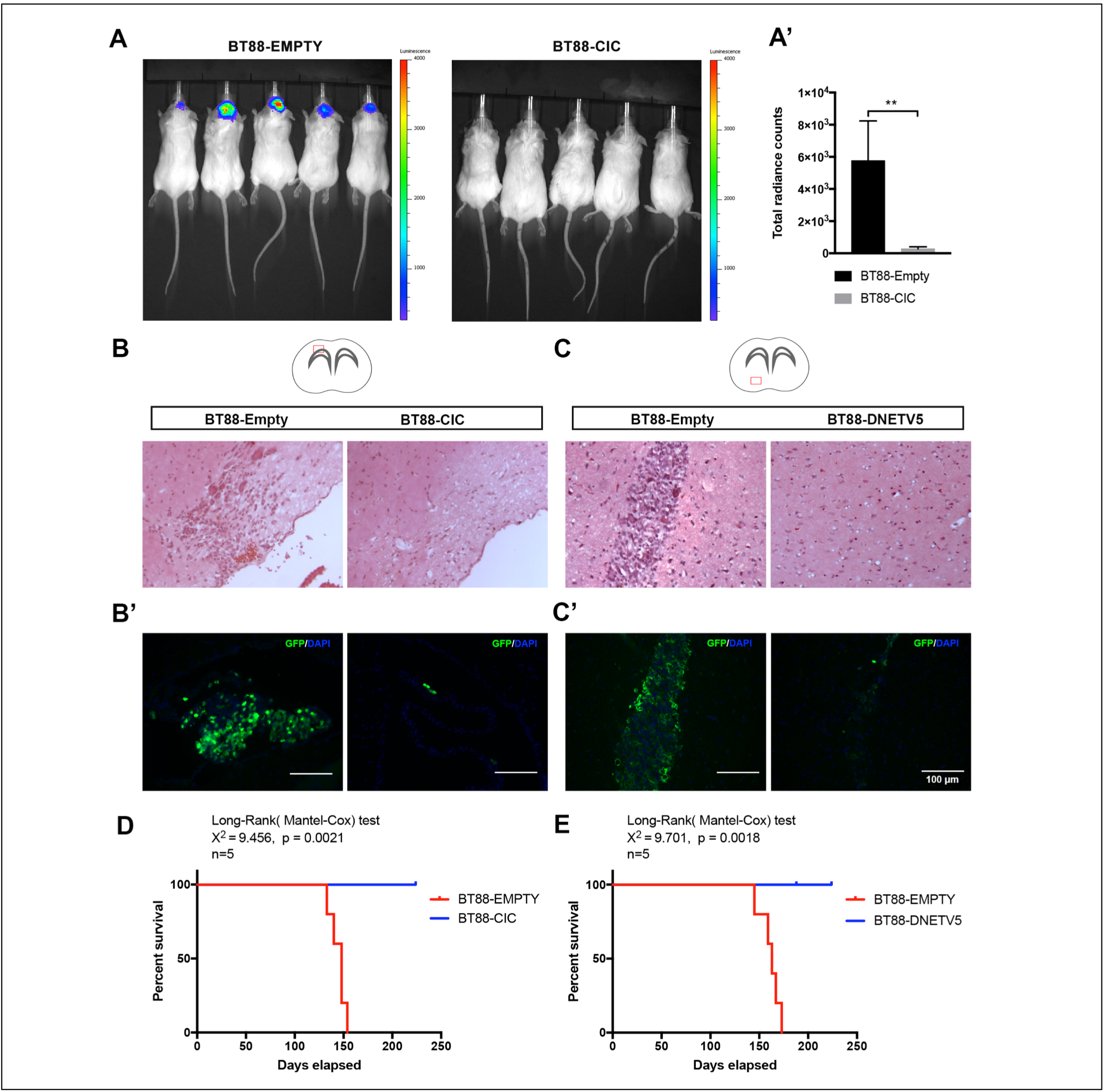
Tumorigenicity of ODG cells is reduced by CIC re-expression or ETV5 blockade. (A, A’) Bioluminescence imaging of NOD-SCID mice at 6 weeks post-implantation of BT88 oligodendroglioma cells stably transfected with either empty-GFP-luciferase or CIC-GFP-luciferase expression vectors. Data shown as mean ± SD from 8 mice per group. Statistical analyses by student’s t test. **p<0.01. (B,B’) Representative images of brain sections from mice implanted with control-GFP-luciferase BT88 cells or CIC-GFP-luciferase BT88 cells stained with H&E or GFP. (C,C’) Representative images of brain sections from mice implanted with control-GFP BT88 cells or DNETV5-GFP expressing BT88 cells stained with H&E or GFP. (D) Kaplan-Meier survival analysis of mice implanted with BT88 cell transfected with empty-GFP-luciferase or CIC-GFP-luciferase. (E) Kaplan-Meier survival analysis for BT88 cell transfected with empty-GFP or DNETV5-GFP.

A similar finding was observed when we assessed the ODG cell lines’ ability to differentiate in response to oligodendrocyte differentiation conditions. Both CIC re-expression and DNETV5 similarly resulted in increased expression of CNPase in the ODG cells compared to respective control cells, indicating at least some improvement in capacity for the tumour cells to differentiate and mature (Figure 9F,F’ bottom row panels; Suppl Fig S7). Interestingly, however, effects on neuronal and astrocytic differentiation, were mixed (Figure 9F,F’ top and middle row panels). Cic re-expression resulted increased the ability of cells to respond to extrinsic neuronal differentiation cues by increasing expressing bIII-Tubulin (TUJ1) and extending neurites; however this was not phenocopied by DNETV5. With respect to expression of GFAP in response to astrocytic differentiation cues, no significant differences were detected with either CIC re-expression or DNETV5 introduction compared to BT-88 control. The latter data suggest that other genetic (or epigenetic) alterations may be more central to driving or maintaining some specific aspects of the tumour cells’ phenotypes, at least in our cultured system. The effects on differentiation in vivo were not evaluated, however, as the lesions resulting from implantation of CIC- or DNETV5-expressing BT88 cells were too small to provide meaningful cell numbers for assessment.

Taken together, our results indicate that Cic regulates NSC proliferation and cell fate in neurodevelopment and oligodendroglioma, and that the pro-proliferative and pro-OPC phenotypes observed with Cic loss are largely mediated through Ets transcriptional de-repression.

## DISCUSSION

Genomic analyses of brain cancers have suggested that *CIC* functions as a tumor suppressor gene in diffuse gliomas, particularly ODGs (3, 4, 36). Yet, to date, knowledge of *CIC*’s role in the brain has been limited. Here we report previously undescribed cell type specific differences in CIC expression in the brain. Moreover, our functional work examining *Cic* function in forebrain development and in oligodendroglioma cells provides new insight into the roles of this putative tumor suppressor in regulating the developmental fate of neural stem/progenitor cells.

We found that genetic ablation of Cic biases cells neural stem cells away from neuronal lineage specification to the selection of glial lineages. Adding to knowledge that RAS/MAPK pathway signaling regulates the switch from neurogenesis to gliogenesis, with *ETV5* implicated in mediating gliogenic competence (17, 34), our results firmly place *Cic* at the intersection of RAS pathway signaling and *Etv5* in the brain, providing a missing link between extrinsic differentiation signals and the execution of transcriptional programs critical for normal neuro- and glio-genesis. With respect to Cic and the anti-neuronal bias that we observed, recent work by Lu, et al. found that disrupting Ataxin-1 (Atxn1)-Cic complexes resulted in abnormal maturation and maintenance of upper-layer cortical neurons – with concomitant effect on behavior, learning, and memory (14). It is unknown, however, the extent to which the effects could be due to specific Ataxin-Cic interactions versus Cic abnormality alone. Our studies, which primarily focused on early lineage selection events, did not systematically address questions of cell maturation (although some of our in vitro morphological findings are consistent with a neuronal maturation defect). Future work would be needed to define the complex neuronal dependencies on Cic, not only in the forebrain but also in the cerebellum where Cic pathology has also been implicated (37) (38).

With respect to neural stem cell proliferation, glial fates, and the implications for ODG; our findings build on the recent observation by Yang, et al. that CIC deficiency increases a population of proliferating OPCs in the brain (13). Our work is not only consistent, but now provides a mechanistic foundation for the observations with identification of symmetric cell division alterations, dissection of lineage selection effects and identification of a downstream mediator. Together, they provide tangible links between *CIC* loss and ODG biology – as both dysregulated proliferation and the persistence of an immature OPC phenotype are cardinal features of this cancer. Many parallels exist between normal OPCs and the cells comprising ODG. In common, both OPCs and ODGs express PDGF, PDGFR, and NG2, which control OPC differentiation (39, 40). Together, the results may thus explain some of these phenotypic features of ODG, and consistent with other reports suggesting that dysregulation of OPCs or OPC-like cells are amongst the early changes in experimental models of gliomagenesis (2, 41).

There are some notable differences between the work reported by Yang, et al. (13), however, and our studies. One difference is that the prior study used HOG cells that, despite their historic name, do not carry the ODG-defining genetic features of 1p19q loss or *CIC* mutation; similarly the Pdgfra-amplified/overexpressed mouse glioma model used is more akin to an RTK-driven GBM than it is to ODG. In deploying the BT-88 and BT-54 models (which genetically and phenotypically faithfully recapitulate ODG), our work may be closer to clinical relevance for ODG. Another difference is that our studies also further clarify the early neuronal-glial fate decision effects of NSCs, with the bias away from neuronal fates and toward glial fates in the setting of CIC loss. Most importantly, however, we now identify *Ets* de-repression as a key mechanism underlying much of the phenotypic effects of Cic loss in NSCs.

Our observation that *DN-Etv5* alone can abrogate so much of the pro-proliferative and pro-OPC phenotype resulting from *Cic* loss was somewhat unexpected. In support of a role specifically for Etv5, our observation that it is the most upregulated factor upon Cic loss, our Etv5 overexpression studies (in vivo and in vitro), and our in vitro Etv5 siRNA experiments all lend independent support pointing to the requirement for this particular Ets factor. However, we do not exclude that other Ets factors such as Etv4 may also contribute. The dominant negative construct may not be entirely specific for inhibiting Etv5 alone; such cross-reactivity has been reported with similar approaches, and further experiments would be required to dissect the relative contribution of different Pea3 factors or other subfamilies of the Ets genes. Nevertheless, the identification of this family of genes as a likely mediator is salient to the development of future therapeutic strategies for ODG. Although it is thought that transcription factors are difficult to target, reports of inhibition of other Ets factors either through peptidomimetic or small molecule approaches (42–44) lend hope that Etv5 could potentially be inhibited for clinical benefit.

Finally, although our findings are relevant to ODG biology, it is recognized that our studies do not directly model the genesis of ODG. Indeed, despite the increase in proliferation observed in Cic-null cells, no tumors were detected when following for up to one year after *Cre* electroporation of the *Cic* floxed mice (data not shown) – in keeping with the concept that several collaborating genetic hits are necessary for oncogenic transformation. Our *Cic* conditional knockout mice, however, will be a useful tool for a future more refined cell type- and stage-specific deletion of *Cic* along with the introduction of other mutations such as of *IDH1*(R132H) (45). We conceptualize *IDH1/2* mutations as a first event in ODG, causing epigenetic changes that that serve to expand the potential pool of cells vulnerable to transformation, with second-hit loss of *CIC* serving to dysregulate proliferation, bias cells to oligodendroglial lineage, and delay them in an immature state. Additional studies using experimental models and human ODG samples are needed to further dissect the relationship of *CIC* mutation to *IDH* mutation. Although one recent study used single-cell RNA-Seq to examine ODG cell subpopulations and did not detect any differences between ODG cells with and without *CIC* mutation aside from the increased expression of PEA3 Ets factors, the number of cells analyzed was limiting for resolution of cell type specific differences, and functional studies had not been performed (46). Similar approaches with larger cell numbers might provide further insight into the biology of ODG with respect to *CIC*.

In summary, we provide evidence that Cic regulates proliferation, fate decisions, and differentiation in neural stem cells, with loss particularly expanding the OPC population. Furthermore, our findings indicate that disruption of the Cic-Etv axis (with abnormal de-repression of *Etv5* downstream of *Cic* loss) is central to the biology of ODG.

## METHODS

### Mice

CD1 outbred mice (Jackson Labs) were used for Cic expression analyses.

*Cic* conditional knockout (*Cic*-CKO) mice were generated at Taconic Artemis by homologous recombination to flank Cic exons 2-11 with loxP sites. The exons targeted are 2-11 of *Cic* short form, transcript variant 1, NM_027882.4 (equivalent to exons 3-12 of the *Cic* long form, NM_001302811.1, Cic transcript variant 4). The targeting vector was generated using clones from the C57BL/6J RPCIB-731 BAC library, and consisted of a 4.0 kb 5’ flanking arm, neomycin resistance cassette (flanked by FRT sites), 5.5 kb loxP-flanked region, a puromycin resistance cassette (flanked by F3 sites), and 6.0 kb 3’ flanking arm. After homologous recombination in C57BL/6N Tac ES cells, generation of chimeric animals, and germline transmission, Neo^R^ and Puro^R^ cassettes were removed via Flp recombination by breeding with a Flp deleter line. The final CIC-CKO allele carries loxP sites in introns 1 and 11, and single residual FRT and F3 sites. Expression of Cre recombinase results in deletion of exons 2-11 and frameshifting of the remaining CIC-S exons 12-20. Genotyping primers were as follows: CIC 5’ flanking region (192 bp wt, 348 bp CKO allele) 5’- AGG AGG TTG TTA CTC GCT ATG G -3’ (forward) and 5’- CTG ATG TCC TAA GAC CTT TAC AAG G -3’ (reverse); CIC 3’ flanking region (273 bp wt, 410 bp CKO) 5’ – CTG TGT CAC TGT CTG CCT TCC -3’ (forward) and 5’ – TGG GTA ATA CCA CCG TGC C – 3 (reverse)’.

*FoxG1-cre* mice (Jackson Labs) were bred to *Cic*-CKO mice to generate telencephalic CIC knockouts and littermate controls. After euthanasia, dissected brains were processed for histology, western blotting, or cell culture. Knockout in tissue or cells post-cre was confirmed by Q-RT-PCR using primer/probe sets for CIC (Applied Biosystems) and by Western blotting with anti-Cic.

In all mouse experiments, the morning of vaginal plug was designated embryonic day 0.5 (E0.5). Both males and females were used.

### In utero electroporation

In utero electroporation was performed as described previously [1] using the following plasmids: pLKO.1-Cic shRNA (Sigma, TRCN0000304642; 5’-CCG GAG CGG GAG AAG GAC CAT ATT CCT CGA GGA ATA TGG TCC TTC TCC CGC TTT TTT G-3’), pLKO.1-non-targeting shRNA (Sigma; 5’- CCT AAG GTT AAG TCG CCC TCG CTC GAG CGA GGG CGA CTT AAC CTT AGG -3’), pCIG2-Cre (which contains Cre-IRES-GFP), pCIC-ETV5 (which contains Etv5-IRES-mCherry), Super piggyBac Transposase (Systems Biosciences, SBI), and piggyBac cargo vector PB513B-1 (SBI) into which cDNAs were cloned for Turbo-Cre and Etv5. The DN-ETV5 consists of an *Etv5-EnR* fusion (gift of Dr. Carol Schuurmans) cloned into the piggyBac construct modified to contain the CAG promoter and GFP-luciferase. DNA was prepared with Endo-free DNA kit (Qiagen) and was injected at 1.5 µg/µl into the telencephalic vesicles of embryos in time-staged pregnant females using a Femtojet 4i microinjector (Eppendorf) then followed by electrical pulses (6 × 43 V, 950 ms interval) applied by platinum tweezer-style electrodes (7 mm, Protech) using a BTX square wave generator (Harvard Apparatus). Post-procedure, embryos were allowed to develop until the time of harvesting. EdU (50 mg/ml in PBS) was injected intraperitoneally into the pregnant dam 30 minutes prior to euthanasia.

### Neural stem cell culture

The VZ of E15 brains were dissected, and tissue was dissociated to single cells using Accumax (EMD Millipore). Cells were grown at 37°C, 5% CO2 in low-adhesion tissue culture flasks (Sarstedt) in mouse neural stem cell (mNSC) media consisting of NeuroCult Proliferation media (Stem Cell Technologies) supplemented with heparin, epidermal growth factor (EGF 20 ng/mL; Peprotech), and fibroblast growth factor (FGF 20 ng/mL; Peprotech). When spheres reached 100–200 µm in diameter, cells were split using Accumax and re-plated at 20,000 cells/mL.

### Transfection of cultured cells

1-4 × 10^6^ dissociated cells were re-suspended in 100 µL of Amaxa Mouse NSC Nucleofector Solution (VPG-1004, Lonza) with 5 µg of plasmid DNA. Nucleofection was performed with a Nucleofector II Device (Amaxa) using the A-033 program. Cells were returned to mouse neural stem cell media for further expansion/selection. For siRNA-mediated Etv5 knockdown, cells were transfected with 50 nM Etv5 or Etv4 ON-TARGET Plus SMARTpool siRNAs (Dharmacon) using Lipofectamine 3000 reagent per manufacturer’s protocol. Cells were assayed 48hrs hours post-transfection.

### Trypan Assay

150,000 cells were seeded per T25 flask in mNSC media and counted after 72 hours on a TC20 cell counter (Biorad) using Trypan blue stain (Thermo). Both dead and live cells were counted.

### Paired cell assay

Dissociated cells were plated at 1,000 cells/ml in mNSC media on dishes coated with CTS CELLstart (Thermo). After 20 hrs, cells were fixed in 4% PFA and immunostained for Ki67. Pairs were scored as symmetric proliferative if both daughter nuclei were Ki67+, symmetric differentiative if both were Ki67−, and asymmetric if one nucleus was Ki67+ and the other Ki67−.

### Neural Colony-Forming Cell Assay

Cells in semi-solid media were prepared using the Neurocult NCFC Assay Kit (Stem Cell Technologies) per manufacturer’s protocol. Cells were plated at a density of 1,650 cells/ml using 1.5 ml per 35 mm culture dish. Dishes were replenished after 7 days with 60 µl of neural stem cell proliferation media supplemented with heparin, EGF, and FGF. Sphere number and size were scored using a gridded scoring dish (Stem Cell Technologies). 8-cell aggregates were used as the cut-off for scoring.

### Lineage-directed differentiation

#### Neuronal Differentiation

NSCs were seeded in mNSC proliferation media (Stem Cell Technologies) on coverslips coated with Poly-L-Ornitihine and Laminin. After 24 hrs, media was replaced with Neurobasal Media, 2% B-27, 2 mM GlutaMAX-I (Thermo). After 4 more days dibutyryl cAMP (Sigma) was added daily to a final concentration of 0.5 mM.

#### Oligodendroglial Differentiation

NSCs were seeded on coverslips coated with Poly-L-Ornitihine and Laminin. After 24 hrs, media was replaced with Neurobasal media, 2% B-27, GlutaMAX-I, 30 ng/mL 3,3’,5-Triiodo-L-thyronine sodium (Sigma).

#### Astrocyte Differentiation

NSCs were seeded on coverslips coated with Geltrex (Thermo). After 24 hrs, media was replaced with DMEM, 1% N2-Supplement, 2 mM GlutaMAX-I, 1% FBS.

### Immunostaining

Tissue was fixed overnight in 4% PFA, cryoprotected in 30% sucrose/PBS, and embedded in Tissue-Tek O.C.T. (Sakura Finetek) prior to cutting 6 µm cryosections on Superfrost Plus slides (VWR). Cultured cells grown on coverslips were fixed for 20 minutes in 4% PFA then rinsed in PBS. Permeabilization was performed with 1× TBST (Tris-buffered saline: 25 mM Tris, 0.14 M NaCl, 0.1% Triton X-100) for 15 min at room temperature (RT). 3% goat or horse serum in TBST x 30 min was used for block. Primary antibodies applied for 1 hr at RT or O/N at 4°C in block. Alexa Fluor secondary antibodies were applied at 1:500 dilution for 1 hour RT. Nuclei were counterstained in 4’,6-diamidino-2-phenylindole (DAPI; Santa Cruz) and mounted with FluorSave Reagent (Calbiochem). For immunostaining of tissue sections when two or more primary antibodies were from the same host species and for comparison of cell-type specific CIC expression, the Opal 4-colour IHC kit (Perkin Elmer) was used per manufacturer’s protocol.

### Western blotting

Cells/tissue was lysed in RIPA buffer (10 mM Tris-Cl pH 8, 1 mM EDTA, 1% Triton X-100, 0.1% sodium deoxycholate, 0.1% SDS, 140 mM NaCl, 1 mM PMSF) with 1X Halt Protease and Phosphatase Inhibitor Cocktail (Thermo). 20 µg protein lysate (50 µg in the case of CIC probing) was run on 4-12% Bis-Tris or 3-8% Tris-Acetate gels (Thermo). Protein was transferred to PVDF membranes in NuPAGE Transfer Buffer (Thermo). Membranes were blocked in TBST with 5% powdered milk. Primary antibodies diluted were applied for 1 hr at room temperature or overnight at 4°C. Horseradish peroxidase-coupled secondary antibodies were applied for 1 hr at room temperature. Membranes were developed using ECL Plus Western Blotting Reagent (GE Healthcare) and X-ray film.

### Transcriptional analysis

Total RNA was extracted using AllPrep DNA/RNA/Protein Mini Kit (Qiagen) per manufacturer’s protocol. For each of 3 biologic replicates per condition, 100 ng RNA was subjected to nanoString analysis on nCounter system (NanoString Technologies) using a custom gene expression codeset containing neurodevelopmental and brain cancer associated genes, and 3 housekeeping genes. Data were analyzed using nSolver software and exported to PRISM software for further statistical analyses. Counts for Etv1, Etv4, and Etv5 were normalized to the average of 3 house-keeping genes (GAPDH, Actin, Tubulin beta chain).

### *In situ* hybridization

Digoxygenin (DIG)-labeled riboprobes were prepared using a 10x DIG-labeling kit (Roche), and ISH was performed as described previously using probes for *GFP* and *Etv5* (17).

### Chromatin immunoprecipitation

50 mg P6 mouse forebrain tissue was use per ChIP assay. ChIP was performed with the SimpleChip Plus Kit with magnetic beads (Cell Signaling) per manufacturer’s protocol with the following exceptions. Tissue was disaggregated using a Dounce homogenizer. In place of micrococcal nuclease, shearing was performed on a UCD Bioruptor on high power with intervals of 30 seconds on, 1 minute off for 30 minutes. 1.5 µl of anti-CIC (PA1-46018, Thermo) was used per IP reaction. Histone H3 antibody and normal rabbit IgG were used as positive and negative controls, as supplied in the kit. PCR primers for the ETV5 promoter region were: 5’-GGTGCAGGCCGAGGCCAGGG-3’ (For) and 5’- CATTGACCAATCAGCACCGG-3’ (Rev).

### Image Analysis

Coronal sections at the level of the anterior commissure were used in all quantitations of cell populations and staining intensities in the dorsal cerebral cortex and corpus callosum. Slides were imaged on AxioObserver fluorescence microscope (Zeiss). Images were processed using CS6 Photoshop software (Adobe) for orientation, false colorization, and overlay/colocalization. Enumeration of cells positive for cytoskeletal, cytoplasmic or membrane proteins was performed manually by counting positive cells using the Photoshop CS6 counting tool. Quantitation of CIC nuclear staining intensity and MBP staining density was performed using FIJI software (47) as follows. Images were first processed using Gaussian blur, then background subtracted. Default threshold limits were used in the threshold tool. Images were converted to binary and watershed was run to separate clumped cells. Nuclear staining was quantitated by using the analysing particles option with separate cut-offs set for each antibody used. For quantitation of MBP expression on tissue sections, FIJI was used by drawing regions of interest on the lateral corpus callosum (cingulum), and the mean integrated density in the regions of interest was calculated.

### Intracranial BTIC xenografts and Bioluminescence imaging

BT88 oligodendroglioma cells were obtained from the Dr. S. Weiss, University of Calgary (33). 1×10^5^ BT88 cells stably transfected with CIC-GFP-Luciferase or DNETV5-GFP or respective empty vectors were stereotactically implanted into the right striata of 6- to 8-week-old NOD/SCID mice. 6 weeks post-implantation, tumour burden was measured either by bioluminescence imaging using IVIS Spectrum In Vivo Imaging System (Xenogen) and/or by histology from euthanized animals. Mice were anaesthetized under isoflurane and intraperitoneal injection of XenoLight D-Luciferin (PerkinElmer) was administered at a dose of 150 mg/kg body weight. Acquisition of bioluminescence images was performed 10 min post-injection. Analysis was performed using Living Image Software by measurement of photon counts (photon/s/cm^2^) with a region of interest drawn around the bioluminescence signal.

### Statistics

Data are represented as mean ± SD from at least 3 biologic replicates for experiments. Comparisons between experimental and control samples were made using 2-tailed t-test or, when there were more than 2 groups, using ANOVA. Tukey’s procedure was applied post-hoc to correct for multiple comparisons during multiple pairwise analyses (e.g. differential Cic expression among cell types). Bonferroni post-hoc correction was applied for statistical analysis of data from NanoString assays. Statistical analyses were performed using Prism sofware (Graphpad).

### Study Approval

Animal use was approved by the University of Calgary Animal Care Committee (protocol AC16-0266) in compliance with the Guidelines of the Canadian Council of Animal Care.

### Data Availability

The data that support the findings in this study are available from the corresponding author upon reasonable request.

### Availability of Biological Materials

Biological materials used in this study are available from the corresponding author upon reasonable request.

## AUTHOR CONTRIBUTIONS

Conceptualization, JAC, STA; Methodology, JAC, CS, STA; Investigation, STA, ADR, RD, LF, MJC, MA, WW, SOL; Acquisition, Analyses and Interpretation of data, JAC, STA; Writing – Original Draft, JAC, STA; Writing – Review & Editing, JAC, SC, JGC, SMR, MAM, WW, RD, SOL; Funding Acquisition, JAC, JGC, CS; Resources, LA, MA, CS; Supervision, JAC, CS, JGC, SMR.

## ACKNOWLEDGEMENTS

This work was funded by Canadian Institutes for Health Research (JAC), Cancer Research Society (CS, JAC), Alberta Cancer Foundation (STA, ADR, LF, JGC), Alberta Innovates Health Solutions (JAC, CS).

