## Supplemental Data File for "Capicua regulates neural stem cell proliferation and lineage specification through control of Ets factors"

### SUPPLEMENTARY FIGURES (S1-S6)

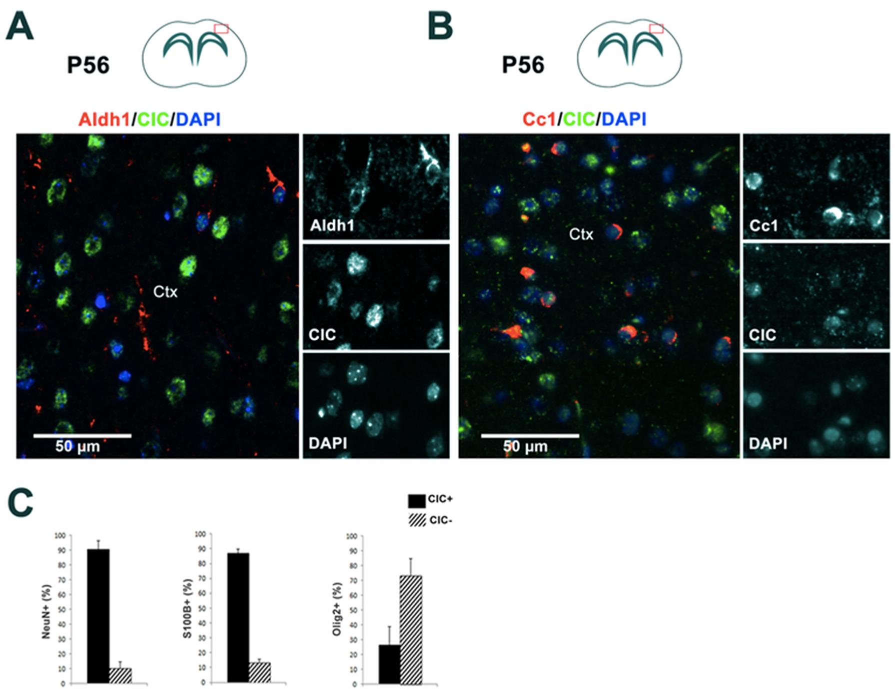


**Figure S1: CIC expression pattern in mammalian neural lineages.** CIC expression in (A) astrocytes labeled by Aldh1 antibody and (B) mature oligodendrocytes labeled by Cc1 antibody. (C) Percentages of NeuN+ (neurons), S100B+ (astrocytes) and Olig2+ (oligodendrocytes) having CIC present or absent, when stained with anti-Cic from Thermo and quantitated as a binary variable using arbitrary threshold. Bars indicate mean±SD. Ctx-Cortex.

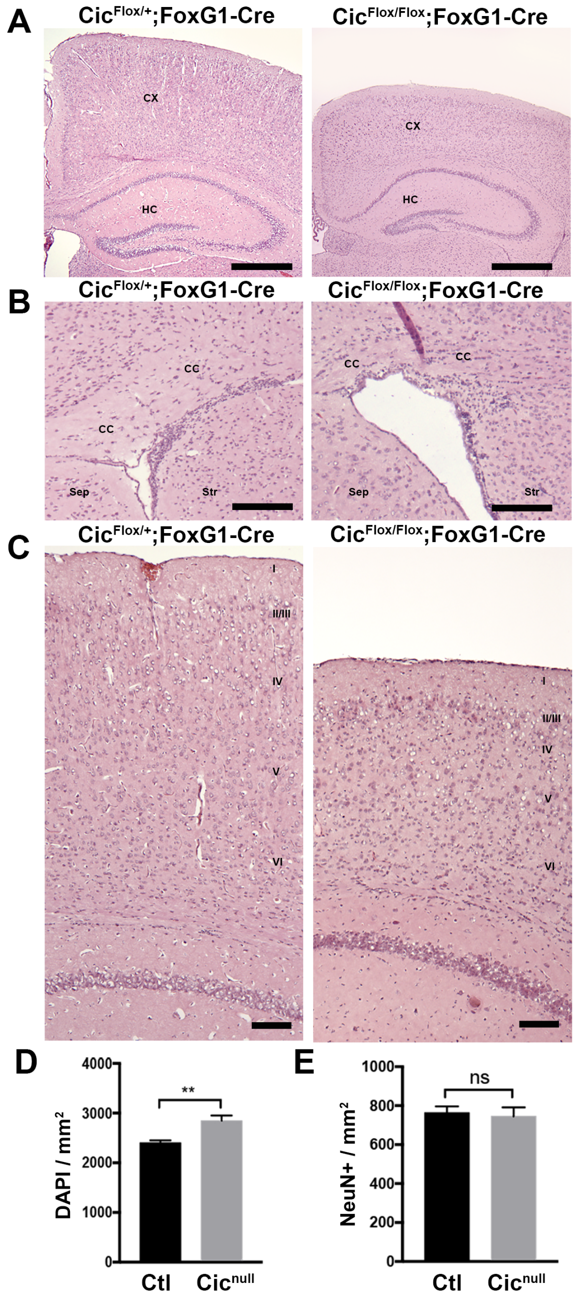


**Figure S2: Additional phenotype of mice with forebrain *Cic* deletion.** Representative images of H&E stained sections of selected areas of mouse brain at P21. Homozygous deletion of CIC results in global decreases in volume of both cortical, white matter, hippocampus, and other deep gray structures. (A) Coronal sections through hippocampus and posterior cerebral cortex at level of the anterior nucleus of the thalamus in mice with CIC heterozygous deletion (Cic$^{Flox/+}$; FoxG1-Cre) or homozygous deletion (Cic$^{Flox/Flox}$; FoxG1-Cre). Scale bar = 500 μm; CX, cortex; HC, hippocampus. (B) Coronal sections through corpus callosum at the level of the anterior commissure. The corpus callosum is thinned with homozygous CIC loss. The subventricular zone, corpus callosum, and adjacent structures also show increased cellularity. Scale bar = 100 μm; CC, corpus callosum; Sep, septal nuclei; Str, striatum. (C) Coronal sections through cortex superior to hippocampus at level of the anterior nucleus of the thalamus show decreased cortical thickness. Scale bar = 100 μm; CX, cortex; cortical layers indicated by I-VI. (D) Cellularity in cortex, as measured by DAPI nuclei per mm$^2$; n=3; ** p <0.01. (E) Neuronal density in cortex, as measured by NeuN+ cells per mm$^2$; n=3; ns, not significant.

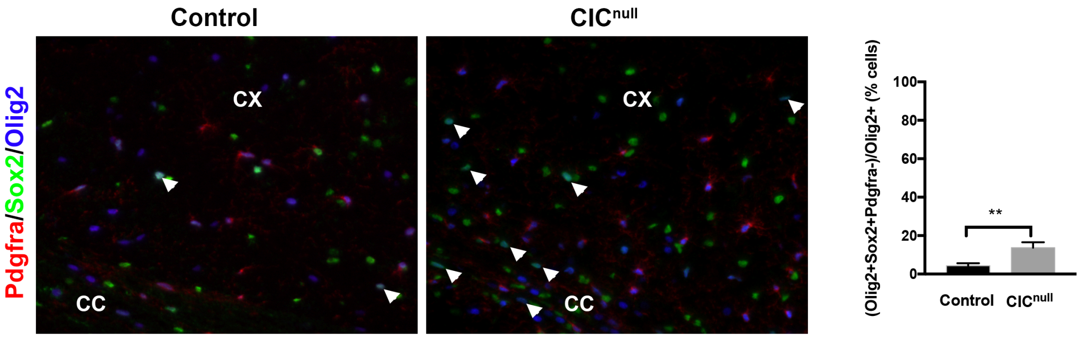


**Figure S3:** (Left panel) Representative images showing Sox2$^{+}$Pdgfra$^{-}$Olig2$^{+}$ cell percentages in control and CIC null mice brains. White arrowheads indicate cells that are Sox2$^{+}$Olig2$^{+}$Pdgfra$^{-}$. (Right panel) Quantitation of Sox2$^{+}$Pdgfra$^{-}$Olig2$^{+}$ cell percentages in control and CIC null brains. (CIC$^{Fl/Fl}$;FoxG1$^{Cre/+}$ 13.67±2.89% vs. Control 4.31±1.33%; n=3, p<0.01). CC, corpus callosum, Cx-cortex.

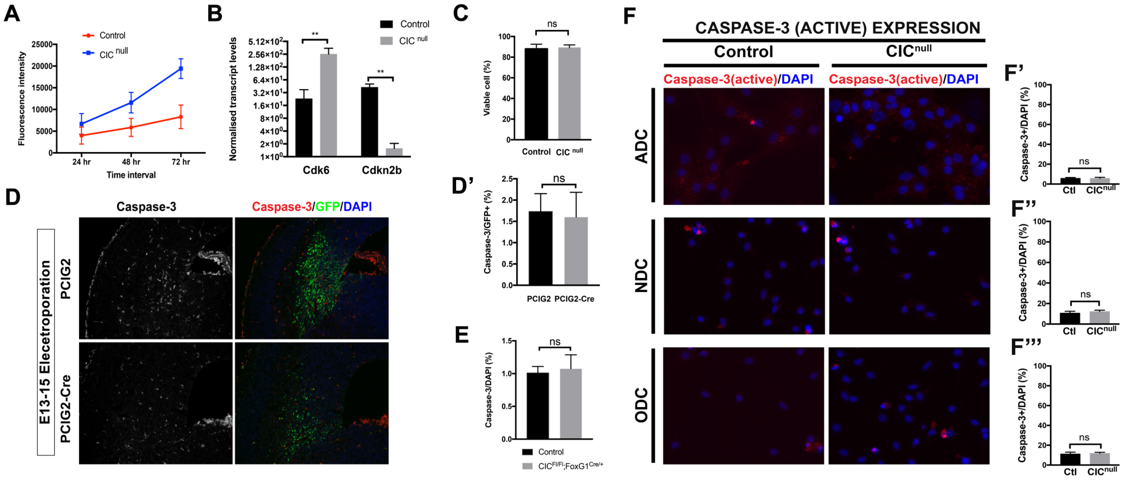


**Figure S4: CIC loss affects proliferation but not apoptosis.** (A) Alamar blue assay showing Cic loss induces higher cell proliferation rates at 24 hrs, 48 hrs and 72 hrs after seeding NSCs at equivalent density. (B) nanoString analysis of Cic null cells and control cell mRNA showing upregulation of *Cdk6* transcripts and downregulation of *P15INK* transcripts by Cic loss. (C) Trypan blue assay shows no difference in the viability of cultured Cic null and control cells when grown in neural stem cell media. (D) Cleaved Caspase-3 immunostaining shows no difference in the apoptotic cell numbers within GFP+ cells between Cic null and control cells. (E) Cleaved Caspase-3 immunostaining shows no difference in the apoptotic cell numbers within GFP+ cells between *Cic$^{Fl/Fl}$;Foxg1$^{Cre/+}$* and control brains. (F) Representative images of caspase-3 immunostaining of all the three neural lineage directed differentiated cells and (F',F'',F''') quantifications showing no significant difference in the apoptosis between control and CIC$^{null}$ cell line. ADC-Astrocytic differentiation condition, NDC-Neuronal differentiation condition, ODC-Oligodendroglial differentiation condition. Bars indicate mean±SD. ns – not significant, ** p<0.01.

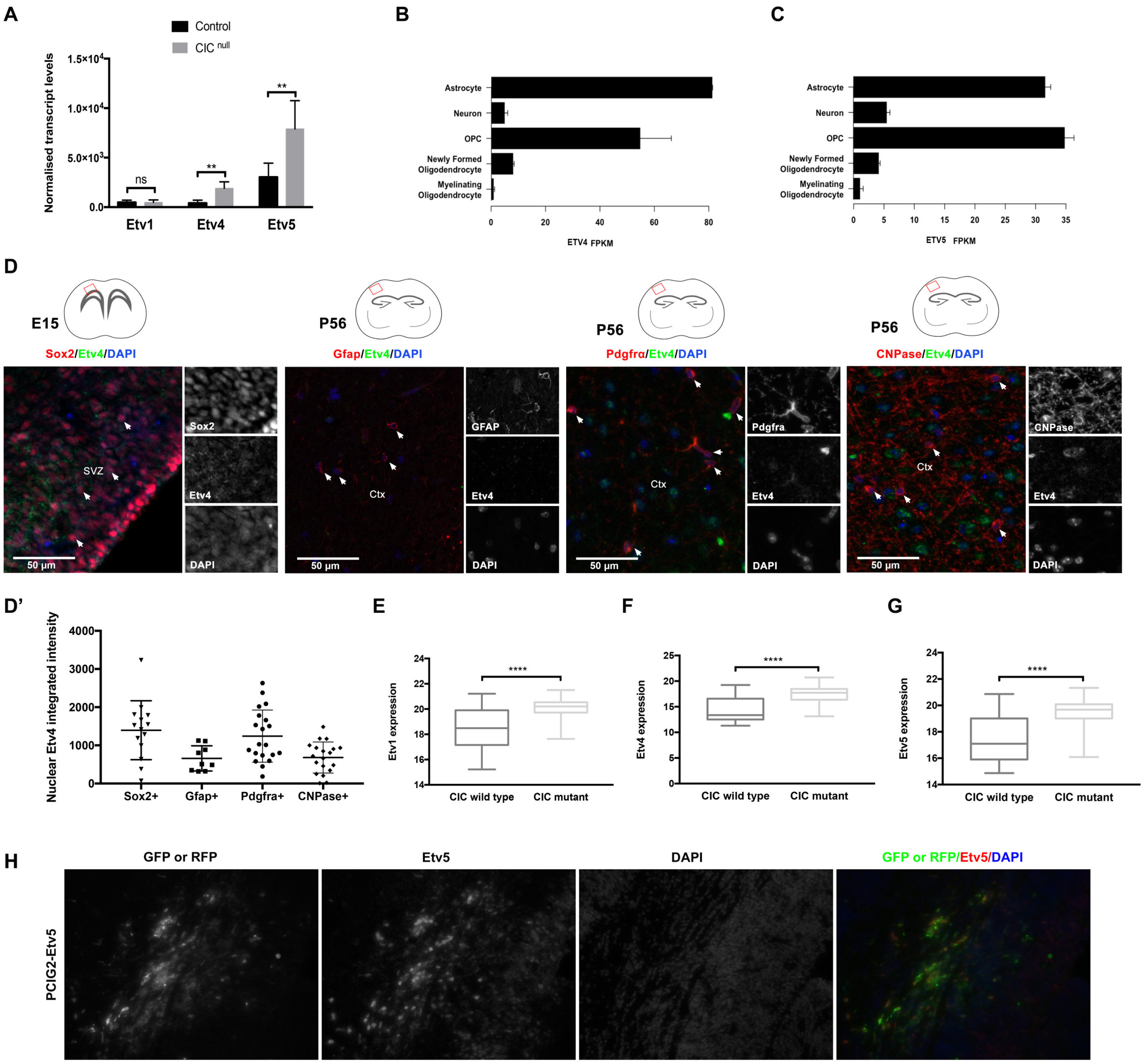


**Figure S5: Inverse relationship of Ets factors with Cic levels.** (A) Transcript levels of *Etv1*, *Etv4* and *Etv5* after 10 days of differentiation of mNSCs. (B) Our data is consistent with Barres lab RNAseq data showing high expression of *Etv4* and *Etv5* transcripts in OPCs and astrocytes (Suppl Ref 1). (C) Etv4 protein expression in mouse brain cortical sections across different neural lineages using specific neural lineage markers. (D,D') Quantification of the Etv4 protein expression across neural lineages. (G) Oligodendroglioma patient cohort (TCGA Low Grade Glioma data set) *ETV1*, *ETV4* and *ETV5* transcripts levels between *CIC*-mutant and -wildtype patients indicates higher expression of *ETS* factors in *CIC*-mutant low grade gliomas compared to *CIC*-wildtype low grade gliomas when controlled for *IDH* mutation status. (H) Immunostaining for Etv5 protein confirms that *in-utero* electroporation of *Etv5* construct into mouse forebrain results in localized increased expression of Etv5. SVZ – Sub-ventricular zone, Ctx- Cortex. ns – not significant, ** p<0.01, **** p<0.0001.

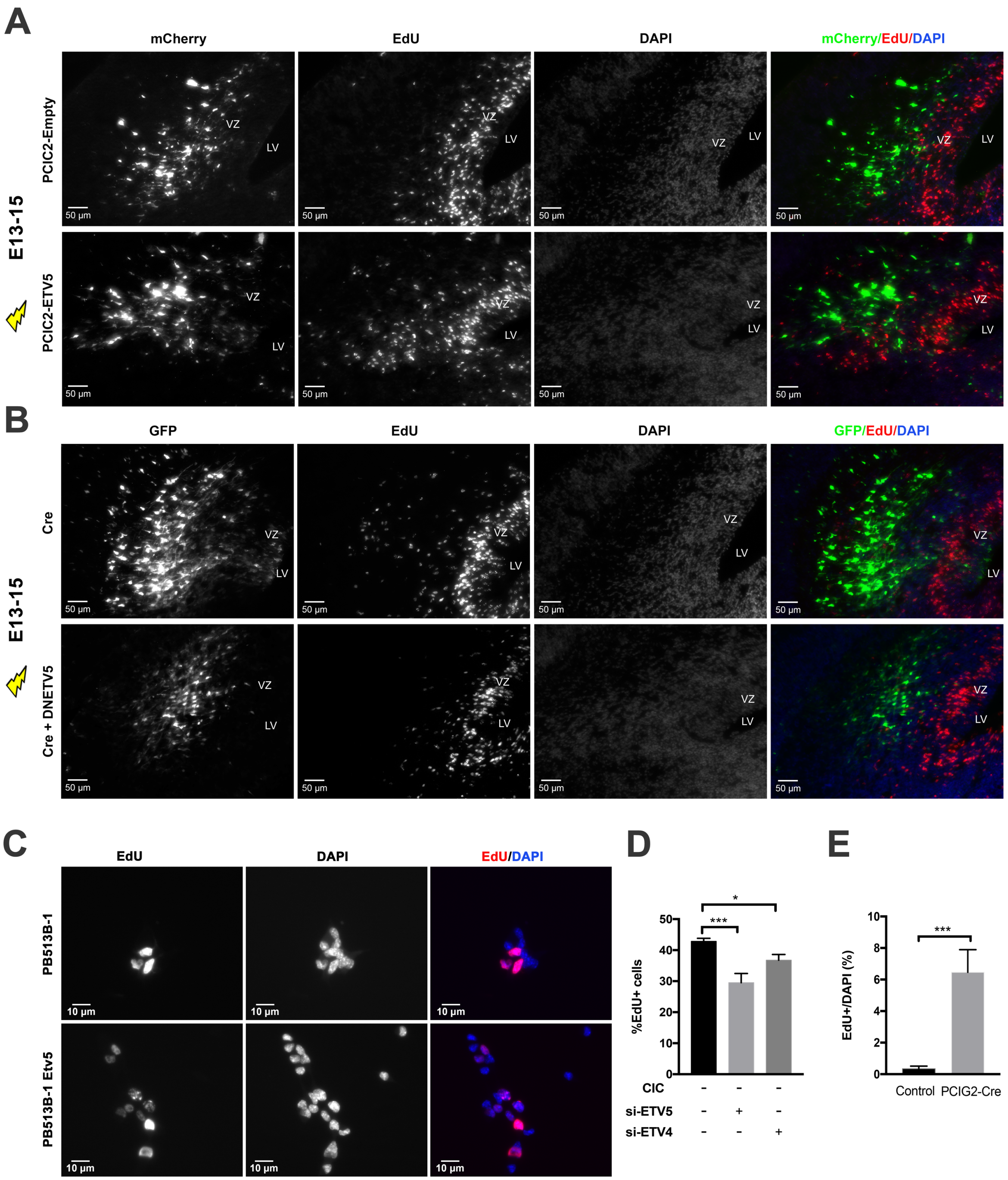


**Figure S6: The proliferative phenotype induced by CIC loss is Etv5-dependent.** (A) Representative images showing Etv5 overexpression increases the percentage of cycling EdU+ cells in electroporated mice with Cre electroporation at E13 and analysis at E15. (B) Etv5 blockade rescues the high proliferation induced by CIC loss. (C) Etv5 overexpression increases percentage of EdU+ cells in cultured mNSCs. (D) Quantitation of percentage of EdU+ cells with siRNA knockdown of Etv4 and Etv5. (E) Quantitation of non-cell autonomous proliferative effects of CIC loss after in utero electroporation of Cre from E13 to E15. VZ – Ventricular zone, LV – Lateral ventricle. * p<0.05, ***p<0.001.

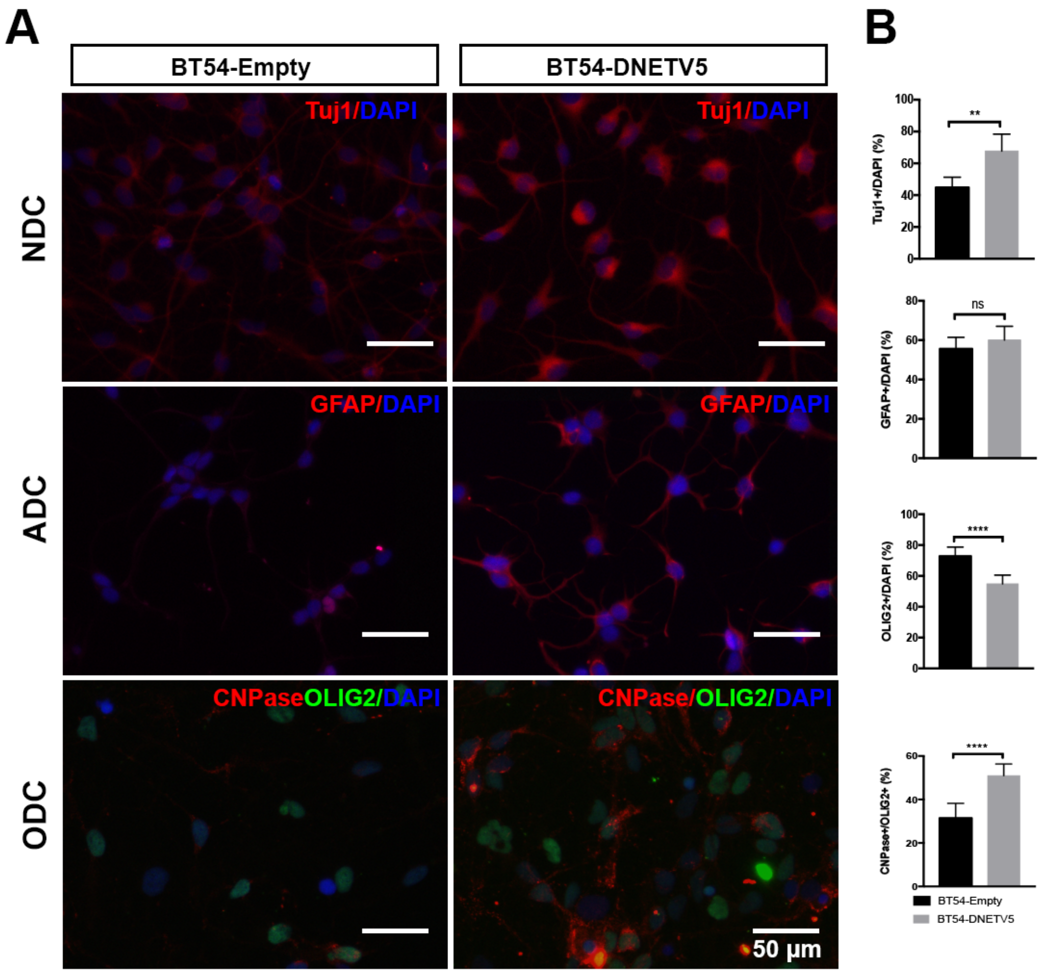


**Figure S7: Etv5 inhibition in the ODG cell line BT-54 increases responsiveness to differentiation cues.** Representative images showing the immunolabelling of BT54-DNETV5 and BT54 control cell lines with neural lineage markers (A). Quantification of the immunostained cells using lineage marker expression. BT54-DNETV5 cells differentiate into neurons with higher efficiency than control BT54 cells under the same differentiation conditions. There was no significant difference in astrocyte numbers under astrocytic differentiation condition between the BT54-DNETV5 and BT54 control line. BT54-DNETV5 cells undergo terminal oligodendrocyte differentiation and become mature oligodendrocyte with higher efficiency than BT54 controls under oligodendrocytic differentiating condition (B). ns – not significant, ** p<0.01, **** p<0.0001.

### SUPPLEMENTARY METHODS

#### The Cancer Genome Atlas data analysis

The Cancer Genome Atlas (TCGA) Lower Grade Glioma (LGG) dataset was downloaded from the online UCSC Xena browser (https://xenabrowser.net/). Tumors that were histologically diagnosed as ODG (grade II or III) or oligoastrocytoma (OA) (grade II or III) that had IDH1 or IDH2 mutations and evidence of 1p19q codeletion (copy number log2(tumor/normal) ≤ -0.5) were included in our analyses. Etv1/4/5 gene expression levels were compared between ODG with wildtype *vs.* mutant CIC.

#### Immunofluorescence quantitation of neural lineages for presence or absence of CIC expression

A fixed number of Neurons (NeuN+), Astrocytes (GFAP+) and Oligodendrocytes (Olig2+) cells in mice brain sections were counted for the presence or absence of CIC expression. The percentages of CIC+ and CIC- cells in each lineage was ploted on a bar graph.

#### Alamar Viability assay

10% Alamar Blue reagent (Life Technologies, DAL1100) was added to cells in 96-well plates and allowed to incubate at 37ºC for 6 hours. The fluorescence reading from each well was then assessed ($\lambda_{ex/em}$: 544 nm/590 nm) using a SpectraMax M2$^{e}$ microplate reader (Molecular Devices).

#### Quantitation of EdU incorporation, non-cell autonomous effects

Equivalent areas of the GFP patch in the IZ (away from VZ/SVZ region) was marked out both in empty construct as well as in Cre electroporated mice brain sections. This area was chosen in order to be able to evaluate similar zones with enough cell numbers in both control electroporation and Cre electroporation conditions. Among GFP- cells in the demarcated zones, EDU+ cells were counted and expressed as a percentage of DAPI stained cells.

#### Antibodies

Primary antibodies directed against the following were used: CIC (1:500 IF, 1:1000 WB; A301-204, Bethyl), CIC (1:100 IF; PA1-46018; Thermo), Olig2 (1:250 IF, 1:1000 WB, MABN50, Millipore), Olig2 (1:500, AB9610MI, Millipore), ETV1 (SAB1403794, Sigma), ETV4 (1:2000 WB, LS-C98380, LS BioSciences), ETV4 (1:500 IF 1:2000 WB, ARP32263_P050, AVIVA Systems Biology), ETV5 (1:1000 WB; sc-22807, Santa Cruz), ETV5 (1:500 IF, 1:2000 WB, 13011-1-AP, ProteinTech), GFP (1:500 IF; A11122, Thermo), GFP (1:250 AF, ab38689, Abcam), SOX2 (1:500 IF, ab92494, Abcam), Turbo-GFP (1:500 IF, OTI2H8, Origene), SOX2 (1:500 IF, 1:1000 WB, #3728, Cell Signalling), SOX9 (1:500 IF, ab76997, Millipore), SOX9 (1:250 IF,1:1000 WB, AB5535, Millipore), PDGFRA (1:500 IF, 1:1000 WB, 3174S, Cell Signalling), PDGFRA (1:250 IF, AF1062, R&D), CC1 (1:500 IF, OP80, Millipore), MBP (1:500 IF, ab40390, Abcam), Nestin (1:500 IF, 1:1000 WB, MAB353, Millipore), Tbr1 (1:500 IF, AB2261, Millipore), Tbr2 (1:500 IF, AB15894, Millipore), Tuj1 (1:500 IF, 1:1000 WB, MAB1195, R&D), GFAP (1:800 IF, 1:1000 WB, MAB360, Millipore), GFAP (1:500 IF, 345860, Calbiochem), ALDH1 (1:250 IF, ab87117, Abcam), NeuN (1:500 IF, MAB377, Millipore), Cre (1:500 IF, 1:2000 WB, NB100-5613, Novus), Ki67 (1:200; ab15580, Abcam), EdU (61135-33-9, Carbosynth US LLC). For routine immunofluorescence, secondary antibodies were: Alexa Fluor-488, -594 and -633) conjugated species-specific antibodies (Thermo) were used at 1:500 dilution. For the OPAL method (OPAL 4-color kit, Perkin Elmer), secondary antibodies used were anti-mouse HRP polymer (DAKO) and anti-rabbit HRP polymer (DAKO).
